# Therapeutic Potential of Neural Rosette Stem Cell-Derived Small Extracellular Vesicles in the Management of Brain Edema

**DOI:** 10.64898/2025.11.30.691418

**Authors:** Signe Emilie Dannulat Frazier, Sofie Krickau Pedersen, Kåre Kryger Vøls, Maria Lie, Carlos Quintana, Frederik Møller Nielsen, Andreas Wrona, Elham Jaberi, Hanni Willenbrock, Qiujia Chen, Xinping Qu, Juan Yang, Nicolaj Strøyer Christophersen, Mie Kristensen, J. Carlos Villaescusa

**Affiliations:** Novo Nordisk, Cell Therapy R&D, DK-2760 Måløv, Denmark; Department of Pharmacy, University of Copenhagen, Copenhagen 2100, Denmark; Novo Nordisk, Novo Nordisk Research Centre China, CN-100193 Beijing, China

**Keywords:** Neural stem cells, Ischemic stroke, edema, Blood brain barrier, neuroprotection, small extracellular vesicles, exosomes

## Abstract

Small extracellular vesicles (sEVs) derived from neural stem cells (NSCs) have gained significant interest for their potential roles in neuroprotection, brain edema management, and the preservation of blood-brain barrier (BBB) integrity. These nanoscale particles facilitate intercellular communication and carry a diverse array of bioactive molecules, including proteins and RNAs, which influence various neurobiological processes. The composition of their cargo depends on the parental cell type, affecting their interactions with recipient cells. In this study, we characterized the cargo of sEVs derived from neural rosette stem cells (NRSCs) at both transcriptomic and proteomic levels and compared it to sEVs derived from cardiomyocyte progenitor cells (CMPCs) and human pluripotent stem cells (hPSCs). Furthermore, we assessed the therapeutic effects of NRSC-derived sEVs in a rat model of transient middle cerebral artery occlusion (tMCAO) to investigate neuronal injury and the progression of brain edema. Our proteomic and miRNA analyses revealed that the sEVs from NRSCs, CMPCs, and hPSCs possess distinct cargo compositions. In an *in vitro* neuronal ischemia model, NRSC-derived sEVs significantly enhanced cell viability, while in an endothelial model, they expedited the restoration of cell barrier integrity. Furthermore, the *in vivo* studies utilizing the rat model of tMCAO showed that intravenous administration of NRSC-derived sEVs at 3, 24 and 48 hours after reperfusion resulted in a significantly faster resolution of brain edema. These findings underscore the influence of the parental cell type on sEV composition and highlight the therapeutic potential of NRSC-derived sEVs in treating brain edema, paving the way for novel therapeutic strategies.

## Introduction

Small extracellular vesicles (sEVs) have garnered increasing attention in neuroscience for their neuroprotective properties and their ability to traverse the blood-brain barrier (BBB), thus presenting a novel avenue for therapeutic interventions in various neurological disorders such as traumatic brain injury, ischemic stroke, and spinal cord injury (Gualerzi *et al*., 2021). These vesicles are currently under investigation as potential treatments due to their regenerative capabilities, neurogenesis promotion and anti-inflammatory effects observed in preclinical stroke models, where intravenous administration has demonstrated improvement in functional outcomes and a reduction in immune response (C. Li *et al*., 2021; Xia *et al*., 2021; Li *et al*., 2023).

The key role of sEVs in mediating neuroprotection can be attributed to their complex cargo, which comprises a range of bioactive molecules, including proteins and microRNAs (miRNAs). These components modulate cellular behaviors and signal transmission in target cells, fundamentally influencing gene expression and promoting cellular resilience against damage (Gregorius *et al*., 2021; Kim *et al*., 2021). Evidence shows that the protein composition and miRNA load of sEVs derived from different cellular sources can significantly impact their therapeutic efficacy, underscoring the importance of cargo characterization in enhancing the neuroprotective functions of sEVs (Zhang, Buller and Chopp, 2019; Barzegar *et al*., 2021; Wu, 2022; Li *et al*., 2025).

While mesenchymal stem cell derived sEVs have historically been the most studied variant, emerging research highlights the superior neuroprotective potential of neural stem cell (NSC)-derived sEVs in stroke models, suggesting that their effectiveness may be linked to shared developmental pathways with neurons or distinct cargo differences (Webb, Kaiser, Scoville, *et al*., 2018; Sun *et al*., 2019; J.-M. Wang *et al*., 2020; Go *et al*., 2021). However, the specific mechanisms underlying these differences remain under investigation, necessitating a deeper exploration into the functional characteristics of sEVs from divergent donor cell types (J. Wang *et al*., 2020).

In the context of acute neurological injury, a series of pathological events unfold in the central nervous system, marked by conditions such as hypoxia, nutrient deprivation, and subsequent cellular waste accumulation, which together compromise BBB integrity and contribute to neuronal apoptosis (Campbell *et al*., 2019; Lv *et al*., 2019; Breitwieser *et al*., 2022). Neurons serve as the primary information-transmitting cells of the brain, characterized by their extensive networks of axons and dendrites, which facilitate the transduction of electrical signals throughout the nervous system (Zeng and Sanes, 2017). These specialized cells rely on a continuous supply of ATP. Any disruption to this flow, such as a lack of blood supply during a stroke, compromises their ability to maintain a transmembrane gradient. This impairment initiates a cascade of events that can lead to anoxic depolarization and subsequent apoptosis (Campbell *et al*., 2019). The significant loss of neurons because of such events contributes substantially to the functional disabilities experienced by patients with neurological injuries. While *in vivo* models have been instrumental in elucidating these pathophysiological interactions, they often focus on specific aspects due to the complexity and variety of readouts needed for comprehensive analyses, thereby underscoring the utility of *in vitro* systems that prioritize specific injury-related phenomena, like BBB dysfunction and edema (Dabrowska *et al*., 2019; Spellicy, 2020; Sawant *et al*., 2022).

The relationship between BBB integrity and brain edema is particularly significant in the aftermath of ischemic strokes, where the stages of cytotoxic and vasogenic edema are intricately linked to BBB disruption (Song *et al*., 2019). The sequence of events initiates with cytotoxic edema resulting from osmotic imbalances, leading to the swelling of cells (astrocytes and endothelial) and culminating in vasogenic edema due to increased BBB permeability (Xia *et al*., 2020; Liu *et al*., 2023; Wu *et al*., 2023). The exacerbation of brain edema is a critical determinant of stroke prognosis, as a high mortality rate is associated with malignant brain edema (Kuang *et al*., 2020; Gu *et al*., 2022).

The therapeutic potential of sEVs in mitigating neurological injuries is promising, but their mechanistic understanding, particularly relating to the influence of their cellular origins and cargo composition, is essential for optimizing their use in clinical applications. In addressing these complex interactions, this study investigates the neuroprotective capabilities of sEVs derived from long-term expandable NRSCs in comparison with those from cardiomyocyte progenitor cells (CMPCs) and human embryonic stem cells (hESCs) sourced from different biological donors. The analysis focuses on characterizing the proteomic and miRNA cargo of these vesicles, thereby elucidating the compositional differences and their implications for protecting endothelial cells against barrier disruption in an animal model of ischemic stroke.

## Results

### Isolation of Small Extracellular Vesicles from Neural Rosette Stem Cell Culture Medium and Polarized Secretion

Neural rosettes are specialized structures that form during central nervous system development, characterized by a radial arrangement of neuroepithelial cells encircling a central lumen enriched with the tight junction protein TJP1 (also known as ZO-1), which is reminiscent of the embryonic neural tube. The formation of neural rosettes involves the establishment of apico-basal cell polarity, wherein TJP1 is localized to the apical surface of these cells, delineating the rosette lumen (Esmailpour and Huang, 2012; Knight *et al*., 2018). In this study, we utilized neural rosette stem cells (NRSCs) maintained at a neural rosette-forming stage, as previously described (Frazier *et al*., 2025). Our objective was to investigate whether this cellular polarization is reflected in the extracellular vesicle (EV) distribution of NRSCs compared to other cell types, such as cardiac multipotent progenitor cells (CMPCs) and human embryonic stem cells (hESCs).

We observed differences in the cellular distribution of the EV marker CD63 among the analyzed cell types (**Figure 1A**). Notably, CD63 accumulated in the lumen of the neural rosettes and co-localized with TJP1 (**Figure 1B**). A similar co-localization pattern was noted for FLOT1 (**Suppl. Figure 1A**), which also co-localized with TJP1 in NRSCs (**Figure 1C**). This finding suggests that the polarized nature of NRSCs may promote a preference for apical secretion of EVs.

**Figure 1.**
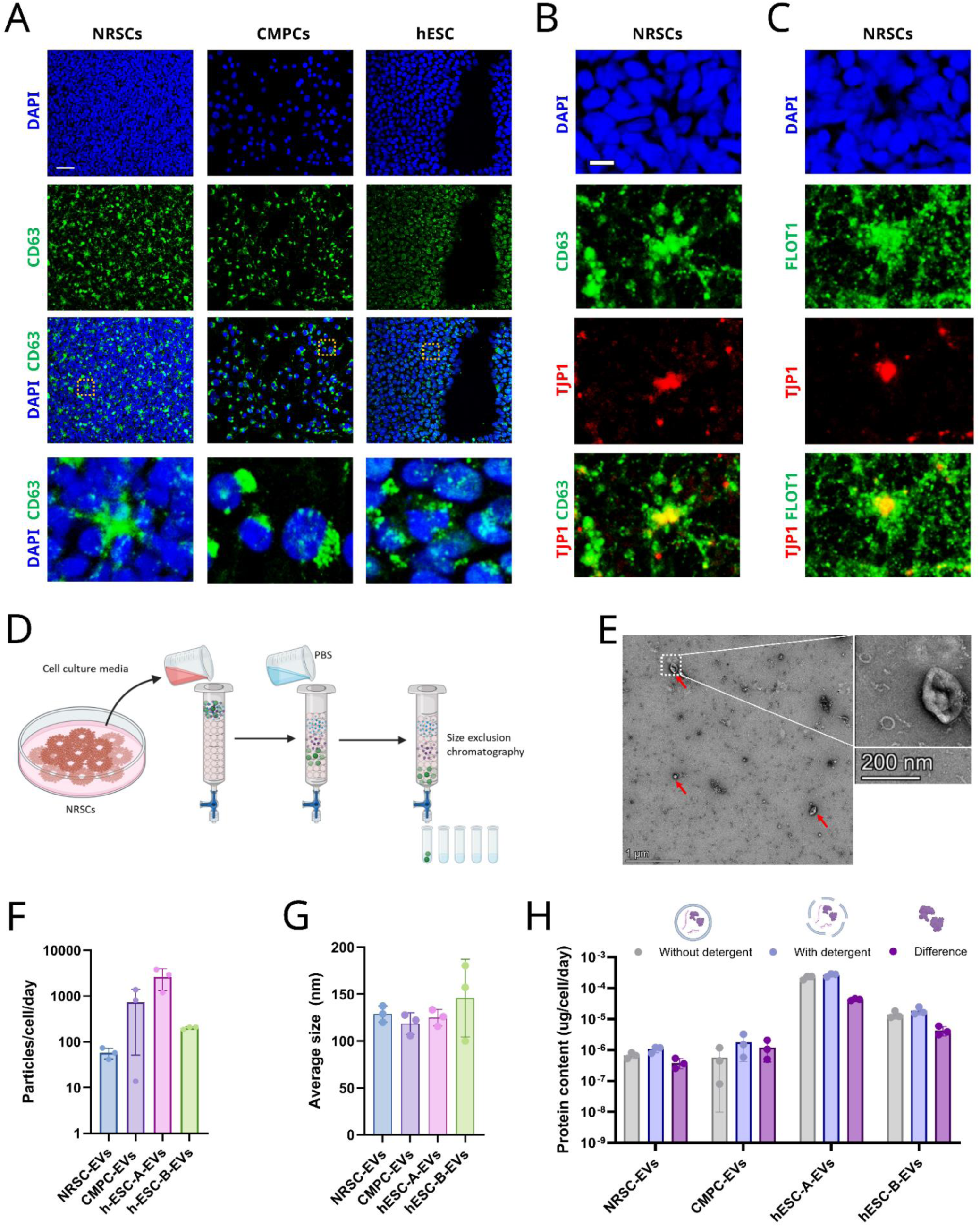
Small extra cellular vesicles can be isolated from NRSC culture medium. **(A)** Representative immunofluorescence images showing the sEV marker CD63 distribution in NRSCs, CMPCs and hESCs. The hESC shown is the hESC-B cell line. scalebar represents 50 µm. **(B)** Immunofluorescence images showing the co-localization of EV marker CD63 and neural rosette lumens, highlighted by TJP1. Scalebar represents 10 µm. **(C)** Immunofluorescence images showing the co-localization of EV marker FLOT1 and neural rosette lumens, highlighted by TJP1. **(D)** Schematic representation of SEC workflow: Cell culture media can be passed through SEC column to isolate sEVs. **(E)** TEM of NRSC-derived EVs showing morphology and size. Scalebar=1µm in the left panel. Scalebar=200µm in the right panel. **(F)** Particle concentration of isolated EVs from 4 different cellular sources. **(G)** The average size of the EVs isolated from 4 different cellular sources. **(H)** Protein concentration of the EVs isolated from 4 different cellular sources. Without detergent represents the proteins located on the exterior of the EV. “With detergent” represents all proteins located both interior and exterior of the sEV and the difference represents the proteins located only on interior of the EV. SEC = Size exclusion chromatography. TEM = Transmission electron microscopy.

To further explore this, we aimed to evaluate the feasibility of isolating EVs from NRSCs and to compare their characteristics with those of EVs isolated from CMPCs and two distinct lines of hESCs (A and B). This comparison sought to uncover differences in EVs derived from various cell types (NRSC, CMPC, hESC) and biological donors (hESC-A and hESC-B). Numerous methods for EV isolation have been described, with ultracentrifugation (UC) being one of the most prevalent. However, gentler methods such as size exclusion chromatography (SEC) may offer advantages in preserving EV integrity and biological activity (Moralès, 2019). Initially, we assessed the ability of UC and SEC to isolate EVs from the culture media of NRSC. Although we successfully isolated EVs using both methods, a direct comparison was not feasible due to significant methodological differences between the techniques and the use of different instrumentation for the measurements (**Suppl. Figure 2A**).

EVs were isolated from concentrated cell culture media of NRSCs, CMPCs, and both hESC lines, followed by separation through a SEC column to fractionate particles based on size (**Figure 1D**). Fractions F1–F5 were combined to form the small extracellular vesicle (sEV) zone, which is defined as containing EVs with sizes below 200 nm. Importantly, the sEV zone includes vesicles that fall within the size range commonly associated with exosomes (**Suppl. Figure 2B, C, D**). Common EV markers, including SYNT1, CD63, CD9, and CD81, were detected and enriched in the pooled F1-F5 fractions compared to the whole cell lysate, confirming EV presence (**Suppl. Figure 2E**). Moreover, transmission electron microscopy was performed to assess the morphology of sEVs isolated from NRSCs, revealing vesicle structures (**Figure 1E**).

The daily particle yield exhibited considerable variability, attributable not only to cell type but also to differing cell culture methods (monolayer versus 3D spheroids) as well as variations in media volume relative to cell count, among other factors (**Figure 1F**). Notably, the size distributions of sEVs derived from all three cell types were broadly similar (**Figure 1G**).

Extracellular vesicles carry protein cargo both on their lipid membranes and within their lumen (Moralès, 2019). To elucidate the relationship between external and internal proteomic cargo, we quantified protein content with and without the use of a detergent. The detergent was applied to solubilize the lipid membrane and extract internal proteins, allowing us to quantify total proteins and exclusively external proteins. The difference between these measures provides an estimation of the internal protein cargo. All cell sources exhibited both internal and external protein cargo, with EVs from both hESC lines showing a higher overall protein content (**Figure 1H**), that could be explained by the increased daily particle yield (**Figure 1G**).

These results indicate that certain EV markers, such as CD63 and FLOT1, accumulate in the lumen of neural rosettes, suggesting an apical secretion preference. Additionally, we found that sEVs can be reliably isolated from NRSC culture medium by UC and SEC and that the collected EVs exhibit classic EV morphology, size distribution and proteomic properties across different cell types.

### Distinct MicroRNA Profiles of Small Extracellular Vesicles from Neural Rosette Stem Cells Compared to Cardiac Multipotent Progenitor Cells and Human Embryonic Stem Cells

We next investigated the transcriptomic profiles of small extracellular vesicles (sEVs) derived from NRSCs in comparison to those from CMPCs and hESCs. sEVs carry a diverse array of nucleic acids, including mRNA, microRNA (miRNA), and non-coding RNAs, which play crucial roles in intercellular communication and the regulation of various biological processes. Analysis of the nucleic acid content revealed distinct profiles across all cell sources (**Suppl. Figure 3A**), prompting us to focus on the miRNA cargo due to its critical involvement in gene expression regulation and its implications in numerous physiological, pathological, and therapeutic contexts (Luarte *et al*., 2016; Zhang, Buller and Chopp, 2019). We profiled the miRNA composition of NRSC-derived EVs and compared it to that of CMPC-derived EVs and those from two different hESC lines (**Figure 2A**), aiming to determine the differences across various cell types and biological donors.

**Figure 2.**
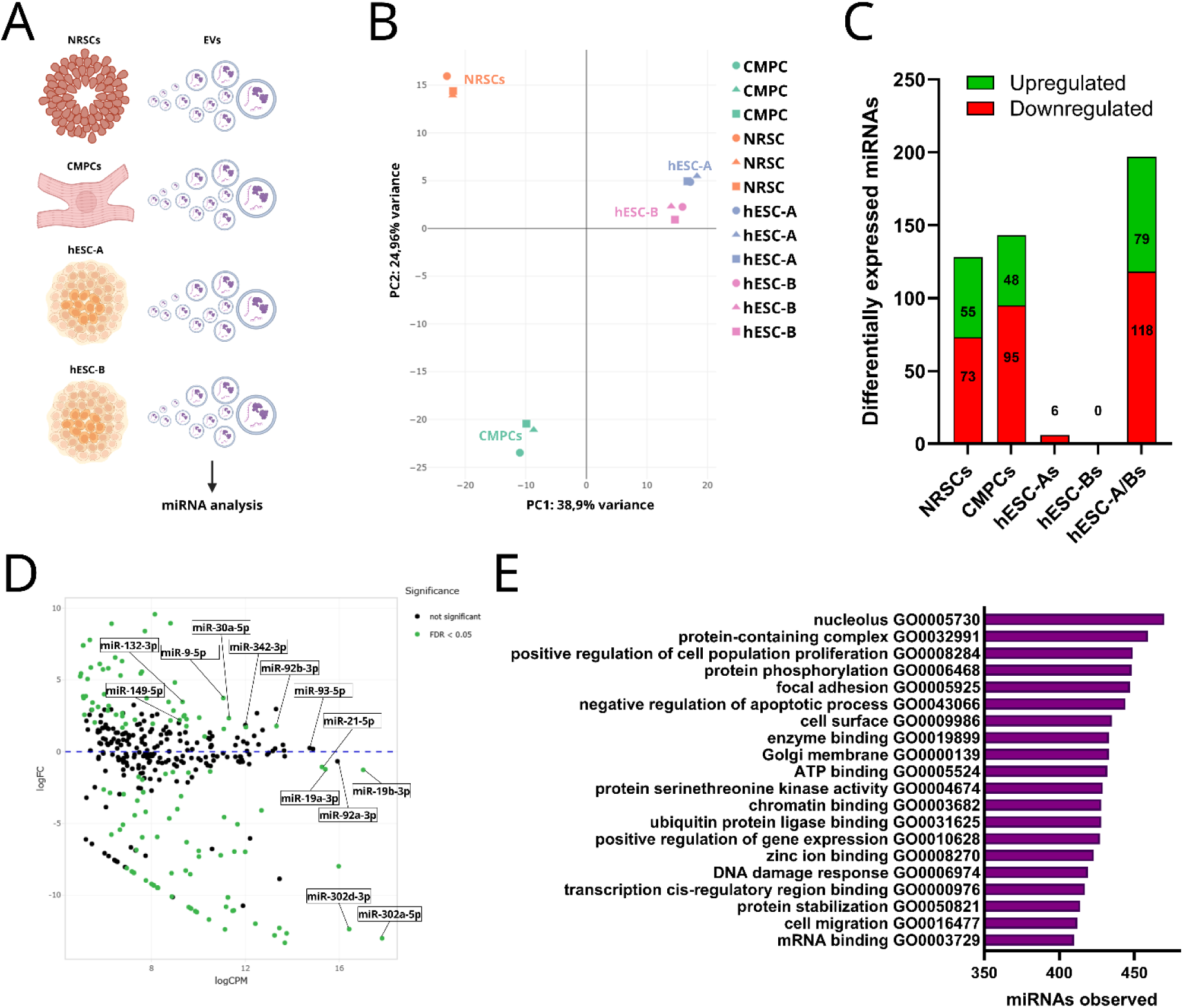
NRSC-EVs carry distinct miRNA cargo. **(A)** Schematic representation of workflow: small EVs were isolated from 4 different cell sources and sequenced to analyze miRNA profiles. **(B)** Principal component analysis (PCA) visualizing the differences in the sEV-miRNA profiles from 4 different sources. **(C)** Number of differentially expressed (DE) miRNAs from each cell-source when compared to all other samples collectively. DE is defined by FDR<0,05. Upregulated genes are shown in green and down regulated genes in red. **(D)** MA plot showing the mean expression counts in relation to the fold change from NRSC-EVs compared to all other cell sources. Significantly differentially expressed miRNAs are shown in green (FDR<0,05). **(E)** Over-representation analysis (ORA) showing the top 20 pathways wherein the NRSC miRNA profile is enriched. The bars represent the number of miRNAs observed in that pathway. N=3 for each cell-source.

While the miRNA profiles of the two hESC lines were similar, notable differences emerged when comparing EVs from different cell types (**Figure 2B, Suppl. Figure 3B**). Specifically, NRSC-derived EVs exhibited greater divergence from hESC-derived EVs, accounting for 38.9% of the variance, compared to their divergence from CMPC-derived EVs. Furthermore, NRSC-derived EVs displayed a unique miRNA cargo pattern, with over 50 upregulated and more than 70 downregulated miRNAs (**Figure 2C**) compared to the other cell sources. CMPC-derived EVs also showed significant upregulation and downregulation of miRNAs compared to other groups. In contrast, the two hESC lines collectively exhibited a substantial number of upregulated and downregulated miRNAs relative to NRSCs and CMPCs, while individual hESC lines showed very few downregulated miRNAs—6 for line A and none for line B—indicating strong similarities between the two lines despite their different donor origins (**Figure 2C**). Notably, neither hESC line contained unique miRNAs in comparison to NRSC, CMPC, or the other hESC-derived EVs, further emphasizing their similarities. In contrast, 19 miRNAs were uniquely expressed in NRSC-derived EVs, and 11 in CMPC-derived EVs, while both hESC lines together exhibited 23 unique miRNAs (**Suppl. Figure 3C**).

Among the uniquely expressed miRNAs in NRSC-EVs were *miR-137-5p*, known for its enrichment in the brain and role in regulating proliferation and promoting neural differentiation (Sun *et al*., 2011; Santos *et al*., 2016), and *miR-98-3p*, which protects endothelial cells from hypoxia-induced apoptosis (Li *et al*., 2015). Although unique, many of these miRNAs were present at low abundances. The most abundant and significantly upregulated miRNAs included *miR-92b-3p*, *miR-342-3p*, *miR-30a-5p*, and *miR-9-5p* (**Figure 2D**). *miR-92b-3p* is associated with promoting neurite outgrowth and facilitating neuronal recovery after injury by negatively regulating PTEN (Chen *et al*., 2019). While less explored in a neurological context, *miR-342-3p* has demonstrated pro-angiogenic properties in cardiovascular disease (Ray *et al*., 2020). Additionally, *miR-30a-5p* is recognized for suppressing inflammatory responses (Fu *et al*., 2018), and *miR-9-5p* is known to promote angiogenesis and enhance neurological recovery following ischemia or traumatic brain injury (Wu *et al*., 2020; Gai *et al*., 2023; Zhao *et al*., 2024).

Interestingly, several significantly upregulated miRNAs, such as *miR-149-5p* and *miR-132-3p*, have been directly linked to the alleviation of edema in rodent stroke models (Wan *et al*., 2018; Zuo *et al*., 2019). The most abundant miRNAs identified from the *miRNA-17∼92* family included *miR-19b-3p*, *miR-92a-3p*, *miR-19a-3p*, and *miR-93-5p* (**Figure 2D**). This family is well-known as regulators of neurogenesis, promoting cell proliferation and inhibiting apoptosis (Xia, Wang and Zheng, 2022). *MiR-302a-5p* and *miR-302d-3p* were also highly abundant in NRSC-derived EVs, as well as in CMPCs and hESC-derived sEVs (**Figure 2D**). *MiR-302a* has been shown to reduce cell proliferation and promote apoptosis by directly targeting VEGFA (Qin *et al*., 2017), while *miR-302d-3p* has been associated with reduced cell viability and increased apoptosis in damaged chondrocytes via ITGB4 targeting, negatively regulating the PI3K-AKT pathway (Sun *et al*., 2021).

Given the distinct miRNA compositions of EVs from different cell types, we sought to elucidate the functional attributes characterizing our NRSC-EVs. To achieve this, we performed an Over-Representation Analysis (ORA) using the miRNA Analysis and Annotation tool (miEAA) (**Figure 2E**). Notably, most miRNAs in NRSC-EVs were found to interact with targets in the nucleolus, the organelle responsible for ribosome production and assembly. Although previous studies have shown the enrichment of ribosomal RNA (rRNA) and specific miRNAs associated with the nucleolus in sEVs (Jenjaroenpun *et al*., 2013; Zhang *et al*., 2021), their precise functions remain to be elucidated. Among the five most enriched biological processes were positive regulation of cell proliferation and negative regulation of apoptosis, which may explain the beneficial effects on cell viability reported in prior studies (Webb, Kaiser, Scoville, *et al*., 2018; W.-Y. Li *et al*., 2021).

Our findings revealed distinct miRNA profiles among the cell types, exhibiting unique miRNAs and a significant number of upregulated and downregulated miRNAs. Notably, the study highlighted the functional implications of the miRNA composition, particularly in processes related to cell proliferation and apoptosis regulation.

### NSC-derived small extracellular vesicles carry distinct proteomic cargo

SEVs carry a diverse array of protein cargo, including neuroprotective factors that modulate inflammatory responses, promote cell survival, and facilitate repair mechanisms in the brain following injury (Zhang, Buller and Chopp, 2019). To elucidate the proteomic cargo of NRSC-derived EVs, we compared it with EVs derived from CMPCs and hESCs (**Figure 3A**).

**Figure 3.**
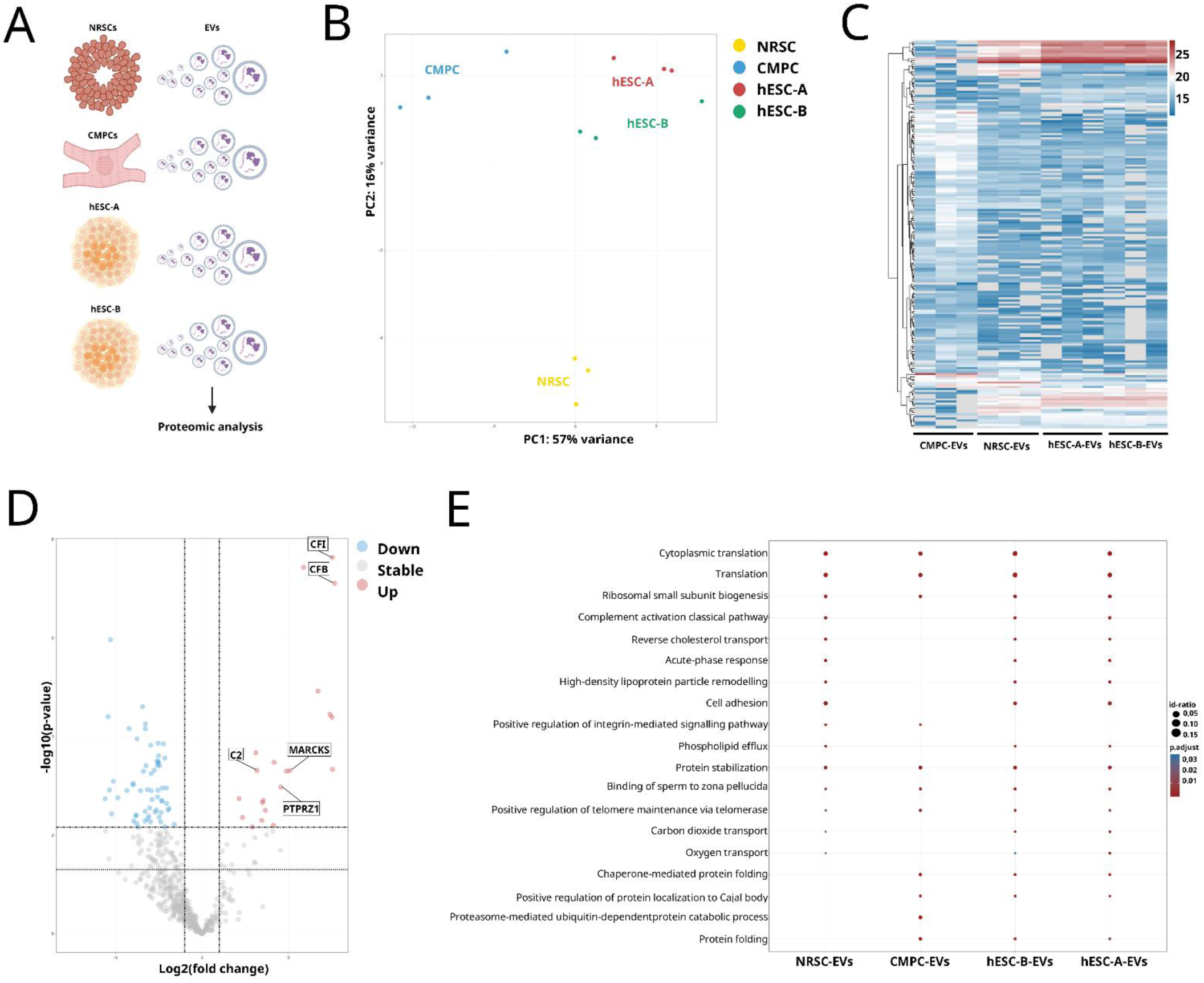
NRSC-EVs carry distinct proteomic cargo. **(A)** Schematic representation of workflow: small EVs isolated from 4 different cell sources and sequence to analyze proteomic profiles. **(B)** Principal component analysis (PCA) visualizing the differences in the proteome in EVs from NRSCs, CMPCs, hESC-As and hESC-Bs. **(C)** Heatmap visualizing the unique patterns of the EV proteome dependent on the cellular source. Heatmap color represents the log2 transformed intensity in individual samples. **(D)** Volcano plot visualizing significantly altered proteins levels in NRSC-EVs compared to all other samples. Red represents significantly increased protein levels; gray represents stable, and blue represents significantly decreased protein levels. **(E)** Over-representation analysis (ORA) showing the top 10 biological processes across all groups. The color of the dots represents adjusted p-value, and the size of the dots represents the number of proteins in that pathway out of the total number of proteins. N=3 for each cell-source.

Our proteomic analysis revealed patterns consistent with the miRNA profile, showing that NRSC-EVs differ more significantly from CMPC-EVs than from hESC-EVs (**Figure 3B**). The proteomic profile of CMPC-EVs was notably distinct compared to that of hESCs (**Figure 3C**). While some soluble proteins may co-purify with sEVs via SEC, particularly in the last F5 fraction (**Suppl. Fig. 2D**), our focus was on proteins which gene expression was validated by scRNA-seq in our NRSCs.

Among the detected proteins in NRSC-EVs, we identified C2, a component of the complement pathway (**Figure 3D, Suppl. Fig. 4B**). The presence of complement protein C2 in sEVs has gained considerable interest due to the multifunctional role of the complement system in immune responses and inflammation (Huang *et al*., 2018). Importantly, the complement pathway is involved in the innate immune response, identifying and lysing pathogens while clearing apoptotic host cells (Mevorach *et al*., 1998; Attali, Gancz and Fishelson, 2004). Efficient clearance of apoptotic cells is particularly relevant in stroke recovery, as it mitigates neuroinflammation and subsequent neurodegeneration (Veerhuis, Nielsen and Tenner, 2011). Additionally, we detected two other components of the complement pathway, CFI and CFB, in NRSC-EVs, however, the number of NRSCs expressing the corresponding mRNA was very low (data not shown).

Another important neuro-specific protein detected in NRSC-EVs was MARCKS (**Figure 3D, Suppl. Fig. 4B**), which is associated with nerve sprouting and axonal regeneration following stroke and spinal cord injury (Carmichael *et al*., 2005; El Amri, Fitzgerald and Schlosser, 2018). Moreover, MARCKS may influence the blood-brain barrier (BBB) by regulating endothelial cell migration and motility (Kalwa and Michel, 2011).

We also detected PTRPZ1 protein, a tyrosine phosphatase receptor type Z, that plays a significant role in the functioning of the central nervous system. It is predominantly expressed in neuronal tissues, particularly in astrocytes and oligodendrocytes, and is involved in essential processes such as oligodendrogenesis, axonogenesis, and neuroinflammation (Baldauf *et al*., 2015; Gascón *et al*., 2024).

We identified 35 proteins unique to NRSC-derived EVs (**Suppl. Figure 4A**). Notably, one of these proteins, SELENOP (also known as SEPP1), has been shown to enhance neural progenitor cell proliferation and neurogenesis when administered in mice (Leiter *et al*., 2022). Additionally, CAT (catalase), an antioxidant enzyme predominantly located in peroxisomes and, to a lesser extent, in mitochondria, decomposes hydrogen peroxide (H₂O₂) into water and molecular oxygen, thereby mitigating oxidative stress. CAT has been demonstrated to protect neurons after injury (Singhal *et al*., 2013), and EVs containing CAT exhibit significant neuroprotective properties against oxidative stress and synaptic damage, effects that are dependent on CAT activity (Bodart-Santos *et al*., 2019).

Several proteins specifically associated with the neuronal niche were also uniquely found in NRSC-EVs, such as THY1 (**Suppl. Figure 4A**), expressed on the surface of neurons and vascular endothelial cells during angiogenesis (Lee *et al*., 1998).

Our analysis also revealed that EVs from NRSCs, CMPCs and hESCs shared a significant association with pathways involved in cytoplasmic translation, mediated by ribosomes (**Figure 3E**). This finding complements our transcriptomic data, which similarly indicated strong associations with ribosome-related processes. Additionally, NRSC-EVs overrepresented proteins associated with the acute-phase response, which is critical for addressing inflammation and injury, as well as promoting cell adhesion—a process closely linked to BBB integrity (**Figure 3E**). Among the cell adhesion-related proteins were several laminins, including LAMA1, LAMA4, LAMB1, LAMB2, and LAMC1, whose absence has been implicated in increased BBB permeability (data not shown) (Bannykh *et al*., 2024; Tremblay *et al*., 2024).

The proteomic analysis of NRSC-derived EVs reveals a complex and unique protein landscape, including neuroprotective factors and components of the complement pathway, which may synergistically influence recipient cells. This intricate composition highlights the potential of NRSC-EVs to modulate inflammatory responses, promote neuroprotection, and support recovery mechanisms following injury.

### Neuroprotective Effects of NRSC-derived sEVs Against Oxygen Deprivation

To evaluate the potential neuroprotective effects of NRSC-derived EVs on ischemic neurons, we established an *in vitro* neuronal oxygen deprivation model. Neurons were differentiated from NRSCs using previously described methods (Frazier *et al*., 2025), and we employed a coupled-enzyme reaction to substantially reduce oxygen levels in the culture medium (**Suppl. Figure 5A**). This approach was inspired by (Baumann *et al*., 2008), where glucose oxidase and catalase were utilized to mimic the hypoxic conditions observed in solid tumors. We tested three different concentrations of the enzymes to determine the efficacy of oxygen depletion (**Suppl. Figure 5B**). It was found that the addition of 3 U/ml glucose oxidase and 180 U/ml catalase effectively reduced oxygen levels to less than 10 mmHg within one minute (**Suppl. Figure 5C**). By supplementing glucose, we extended the ischemic condition for up to six hours (**Suppl. Figure 5D**). Although variable, five hours of oxygen deprivation reduced cellular viability to approximately 50%, providing an opportunity to assess the effects of NRSC-derived EVs (**Suppl. Figure 5E**).

After five hours of oxygen deprivation, NRSC-EVs were added along with EVs derived from CMPCs and hESCs to the neurons (**Figure 4A**) and treated for two days. After that, cellular viability was evaluated employing two different assays based on reducing potential as an indicator of metabolic active cells or on ATP content. The most notable recovery was observed in neurons treated with NRSC-derived EVs, indicating that the effects are likely related to the cell type producing the sEVs (**Figures 4B, C**). Specifically, NRSC-derived EVs enhanced overall cellular viability, as indicated by an increase in reducing potential from 28% to over 50% (**Figure 4B**) and an increase in total ATP levels from 15% to over 30% (**Figure 4C**). While a relatively high recovery was also noted in neurons treated with CMPC-derived EVs, this group exhibited greater variability compared to the NRSC-EV group. Brightfield imaging further corroborated these findings, revealing a higher proportion of visibly healthy neurons following exposure to both NRSC-derived and CMPC-derived EVs (**Figure 4D**).

**Figure 4.**
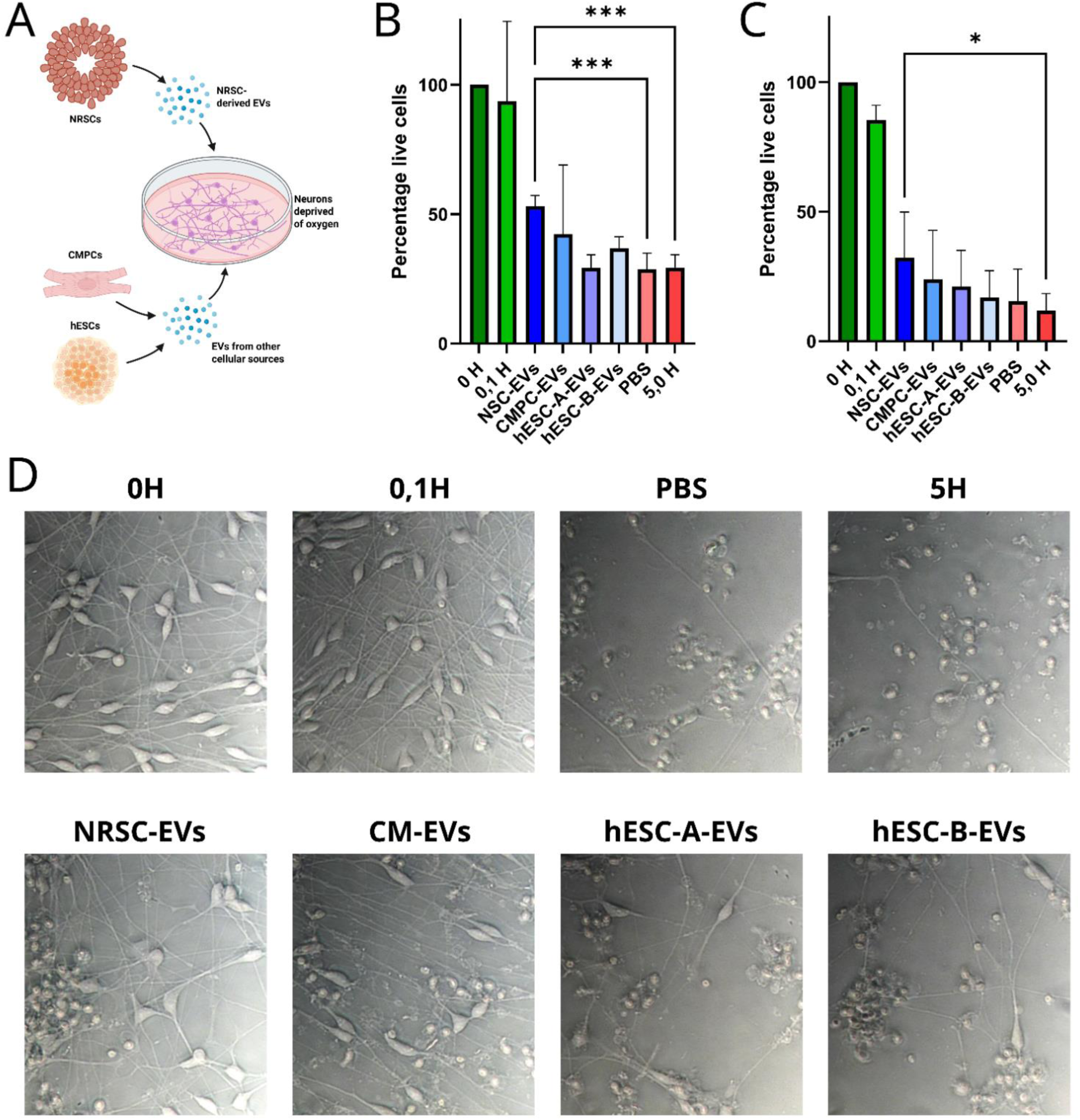
NRSC-EVs rescue neurons deprived of oxygen superior to EVs from other cellular sources. **(A)** Schematic illustration of experimental design. NRSC-EVs were added to oxygen-deprived neurons, and their effect compared to EVs derived from other cell sources. **(B)** Viability measurement based on reducing potential of neurons after 2 days of treatment with EVs from different cellular sources. **(C)** Viability measurement based on total ATP level of neurons after 2 days of treatment with EVs from different cellular sources. 0 H, 0,1 H and 5,0 H all refers to the number of hours that the neurons were deprived of oxygen. All EVs treated groups and PBS were subjected to 5,0 H of oxygen deprivation. Data is shown as mean +/− SD. Statistical test done is two-way ANOVA with post-hoc Dunnetts multiple comparisons test, *=P<0,05 ***=P<0,001. N=3 **(E)** Brightfield images showing the morphological changes in neurons after 2 days of treatment with EVs from different cellular sources. Images are taken in 40X magnification.

To confirm that the observed effects were not influenced by the recipient neuronal subtype, we replicated the experiments using NRSC-EVs and hESC-EVs on neurons differentiated from NRSCs via an alternative protocol (**Suppl. Figures 6A, 6B**). Consistently, we observed similar trends, with NRSC-EVs demonstrating stronger effects on overall cellular viability when compared to hESC-EVs (**Suppl. Figures 6C, 6D, 6E**).

Our findings demonstrate the superior neuroprotective effects of NRSC-derived EVs on ischemic neurons, significantly enhancing cellular viability following oxygen deprivation. These results underscore the potential of NRSC-EVs as a promising therapeutic approach for minimizing neuronal ischemic injuries.

### NRSC-Derived sEVs Promote Cellular Barrier Integrity

Structural alterations and leakage of the blood-brain barrier (BBB) frequently occur because of brain trauma, leading to the formation of brain edema and the entrance of harmful substances into the brain, which can potentially worsen neuronal loss. The critical involvement of the BBB in maintaining neurological integrity has been documented, indicating that its disruption significantly correlates with various forms of brain injuries (Harting *et al*., 2010; Zhang, Chen and Lu, 2015; Shah and Kimberly, 2016). To assess if NRSC-EVs could influence BBB integrity, we used an *in vitro* BBB model of primary mouse brain capillary endothelial cells (BCECs) (**Suppl. Figure 7A and 7B**). The BCECs were harvested from mice cortices and cultured on transwell-filter inserts for 4 days before the endothelium was deprived of basolateral medium to remove nutrients, thereby mimicking stroke-like conditions. Thereafter, the cells were cultured for 3 additional days being subjected to consequential starvation and waste accumulation. After this period, the BCECs were exposed to the NRSC-EVs or PBS as a control group, and the barrier integrity was assessed by Trans-endothelial Electrical Resistance (TEER) measurements and permeability studies (**Figure 5A**). We did observe a gradual recovery of the barrier integrity in both groups, as evident by increase in TEER values, however, the BCECs that were exposed to NRSC-EVs recovered faster and displayed higher resistance values (**Figure 5B**). After 3 days, the TEER values of BCECs exposed to NRSC-EVs resembled that of BCECs that were not deprived of basolateral medium (healthy BCECs) (**Suppl. Figure 7C**).

**Figure 5.**
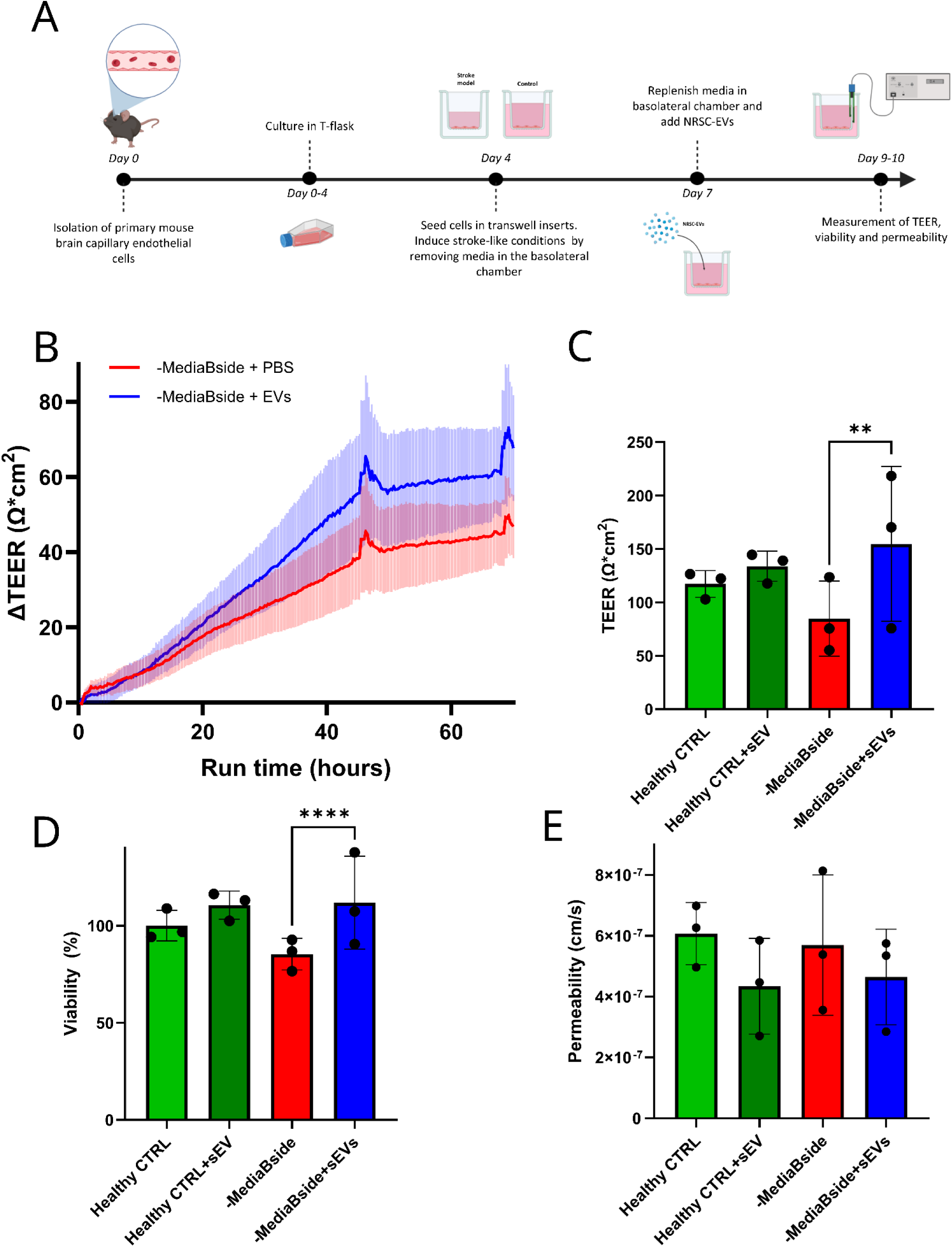
NRSC-EVs promotes faster recovery of injured BBB-model. **(A**) Schematic illustration of experimental design. BCECs were harvested from mice and cultured firstly in T-flasks and then on transwell filters. Stroke-like conditions were induced by removing the medium from the basolateral chamber for 3 days before EV-treatment was initiated. **(B)** ΔTEER (normalization to 0 from initial TEER value) of BCECs over 70 hours after addition of NRSC-EVs or PBS. Data is shown as mean+/− SEM. N=6 **(C)** Endpoint TEER measurement of BCECs after 48 hours of EV treatment. Data shown as mean +/− SD. Statistical test performed is two-way ANOVA with post-hoc Šídák’s multiple comparisons test, **=P<0,01. N=3 **(D)** Viability measurement based on total ATP level of BCECs after 48 hours of EV treatment. Data shown as mean +/− SD. Statistical test performed is two-way ANOVA with post-hoc Šídák’s multiple comparisons test, ****=P<0,0001. N=3 **(E)** Apparent permeability of sodium fluorescein across BCECs after 48 hours EV treatment. Data shown as mean +/− SD. N=3.

To complement our observations, an additional study was conducted using the same experimental setup but utilizing end-point readouts after 48 hours of NRSC-EV exposure. In this setup, similar tendencies were observed in the barrier integrity, with TEER values significantly higher in the BCECs treated with NRSC-EVs (**Figure 5C**). Moreover, cell viability was also significantly increased in the BCECs exposed to NRSC-EVs (**Figure 5D**). Finally, we wanted to determine whether these changes would affect the barrier functionality of the BCECs. To address this, we measured the permeability of sodium fluorescein across the endothelial layer over time. Our data showed a tendency toward slightly reduced transport across BCECs exposed to NRSC-EVs in both the healthy BCECs and the sEV treated groups, although not statistically significant, thus suggesting a mild effect rendering the barrier less permeable (**Figure 5E**).

These results collectively indicate that NRSC-EVs exert a direct and beneficial effect on damaged BCECs, enhancing barrier integrity. This is evidenced by increased TEER values and improved cell viability, which in turn leads to reduced permeability.

### NRSC-EVs Enhance Edema Resolution in Rats Following Experimental Ischemic Stroke

To gain a deeper understanding of the effects of NRSC-EVs on neuronal and endothelial injury, we conducted a study in which rats were subjected to a 90-minute transient middle cerebral artery occlusion (tMCAO) and subsequently administered NRSC-EVs or PBS as a control intravenously at 3, 24, and 48 hours post-reperfusion. The impact of NRSC-EVs was evaluated through MRI and behavioral assessments on days 3, 7, and 28 (**Figure 6A**).

**Figure 6.**
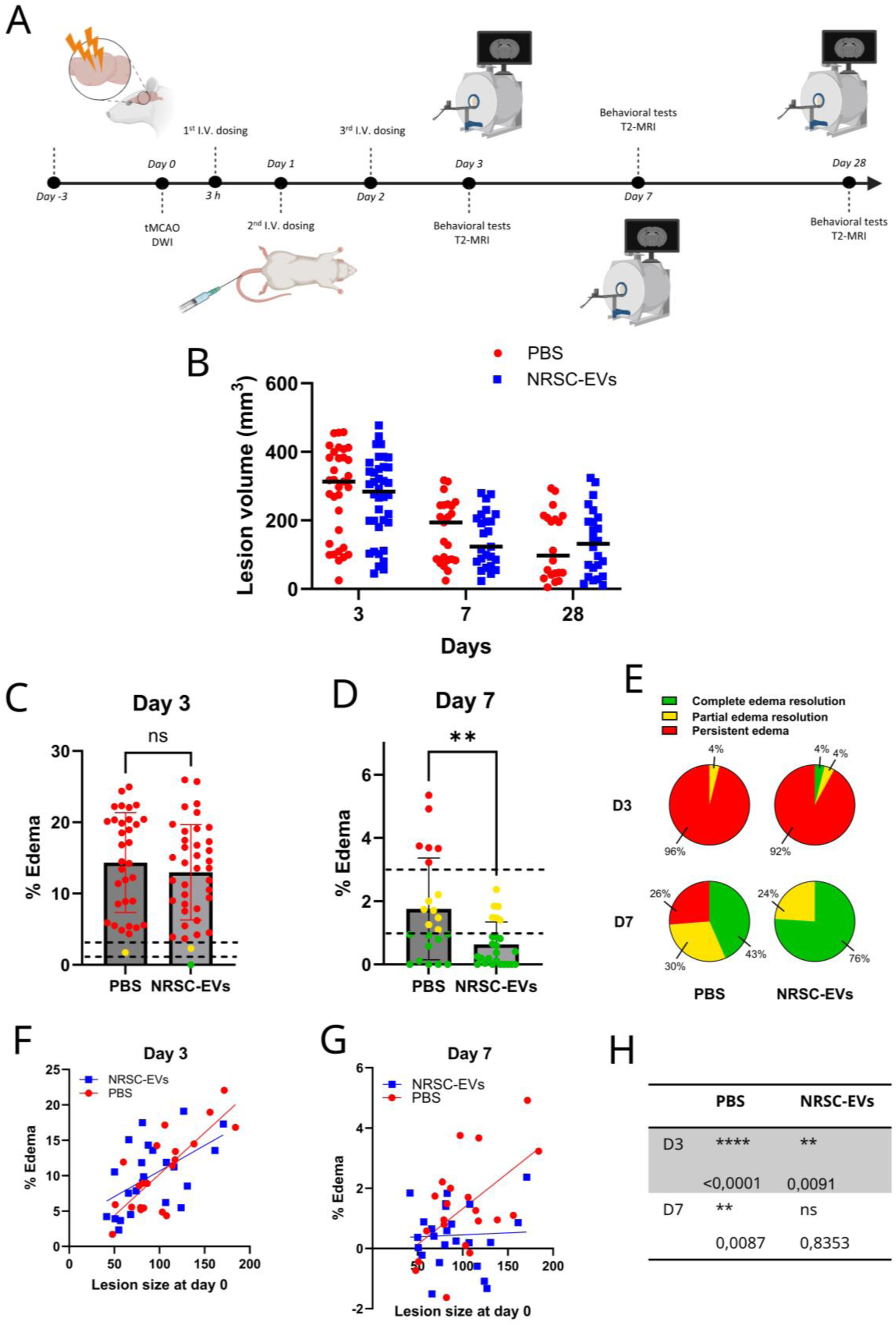
NRSC-EVs facilitates faster alleviation of edema after stroke in rats. **(A)** Schematic illustration of experimental design. Rats were subjected to tMCAO at day 0, dosed with NRSC-EVs after 3 h, 24 h and 48 h and finally the therapeutic outcome was assessed at day 3, 7 and 28. **(B)** Lesion volume (mm^3^) of all animals alive on day 3, 7 and 28. Data shown as median. For NRSC-EV group, N=37 on day 3, N=25 on day 7 and N=24 on day 28. For PBS group, N=33 on day 3, N=23 on day 7 and N=20 on day 28. **(C)** Percentage edema in the ipsilateral hemisphere from all animals that survived until day 3. Data is shown as mean +/− SD. Statistical test done is unpaired t-test. **(D)** Percentage edema in the ipsilateral hemisphere from all animals that survived until day 7. Data is shown as mean +/− SD. Statistical test done is unpaired t-test, **=p<0,01. **(E)** Severity interpretation of % edema values. The percentages represent animals out of the whole group within that severity category. **(F)** Relationship between initial lesion size and % edema on day 3. The lines represent simple linear regression. **(G)** Relationship between initial lesion size and % edema on day 7. The lines represent simple linear regression. **(H)** Test of correlation between initial lesion size and % edema showed a clear correlation on day 7, but only for the PBS group on day 3.

Initially, 96 animals were enrolled in the study. Of the 96, 10 animals were excluded due to insufficient lesion size, 28 due to adherence to strict humane endpoint criteria (body weight loss exceeding 20%), and 12 due to seizure activity. Notably, of the 12 animals that experienced seizures, 8 were from the PBS (control) group, while 4 were from the NRSC-EV group. To ensure successful occlusion and assess the distribution of lesion sizes across groups, diffusion-weighted imaging (DWI) was performed on day 0, revealing no statistically significant differences in lesion size at that time point (data not shown). Furthermore, lesion sizes evaluated by T2-MRI on day 3, 7 and 28 showed no major differences between the NRSC-EVs and PBS groups (**Figure 6B**).

The extent of edema in the ipsilateral hemisphere was measured on day 3 and 7. While no significant differences were observed among the NRSC-EVs and PBS groups on day 3 (**Figure 6C**), a statistically significant resolution of edema was identified in the NRSC-EV group on day 7 compared to the PBS group (**Figure 6D**). Specifically, 76% of the rats in the NRSC-EV group exhibited complete resolution of edema, while only 43% of the rats in the PBS group demonstrated similar recovery. Additionally, while 26% of the rats in the PBS group retained persistent edema by day 7, none of the rats in the NRSC-EV group showed ongoing edema (**Figure 6F**). We also investigated the correlation between edema and the initial lesion size, finding a clear relationship between these parameters in both groups on day 3 (**Figures 6G and I**). However, on day 7, this correlation was absent in the NRSC-EV-treated rats, although a significant correlation persisted in the PBS control group (**Figures 6H and I**). This suggests a faster resolution of brain edema in the NRSC-EV group.

Only minor differences between the NRSC-EV- and PBS-treated rats were noted in the limb placing tests, 20-point neuroscores, and 7-point neuroscores, with a high degree of spontaneous recovery observed in both groups (**Suppl. Figure 8A, 8B and 8C**). The limb placing test assesses the ability of the rats to place their limbs back onto a surface after being pushed away, whereas the neuroscores quantitatively evaluate neurological deficits, with higher scores indicative of better recovery.

This data collectively suggests that NRSC-EVs may promote quicker resolution of edema in rats subjected to tMCAO compared to the control group.

## Discussion

In this study, we evaluated the potential of NRSC-derived small extracellular vesicles (sEVs) as neuroprotective agents in neurological injuries characterized by ischemic hypoxia. Such conditions can lead to breaches in the blood-brain barrier (BBB) and subsequent edema formation, a process that may be relevant across various clinical scenarios, including ischemic stroke and traumatic brain injury.

Our sEVs were derived from high-purity, long-term expandable NRSCs of forebrain identity, as previously documented (Frazier *et al*., 2025). Neural rosettes are pivotal structures in the development and function of neural stem cells, and play a critical role in balancing proliferation and differentiation during neural development (Medelnik *et al*., 2018; Gao *et al*., 2020; Miotto *et al*., 2023). Notably, we observed an apical accumulation of sEVs in our NRSCs, indicating a specialized secretion pattern (**Figure 1A, B, and C**).

Recent research has begun to uncover the roles of sEVs released from NSCs in neurogenesis and neural regeneration. For instance, sEVs derived from NSCs have been shown to enhance neurogenesis and support neuroprotection in models of traumatic brain injury, presenting a novel strategy for therapeutic applications (Abedi *et al*., 2022). Our NRSC-EVs demonstrated multifaceted benefits in both *in vitro* and *in vivo* models. *In vitro*, we observed a protective effect of NRSC-EVs on hypothalamic-like neurons subjected to hypoxic conditions. Following 5 hours of hypoxia, NRSC-EVs significantly increased neuronal viability compared to sEVs from CMPCs and hESCs (**Figure 4**). Furthermore, NRSC-derived sEVs effectively protected neurons from hypoxic conditions, even when derived from a different protocol aimed at achieving a more ventral midbrain neuronal identity (**Suppl. Figure 6**). These findings are consistent with previous studies demonstrating the neuroprotective properties of NSC-derived sEVs (Webb, Kaiser, Scoville, *et al*., 2018; Sun *et al*., 2019).

The critical role of the BBB in maintaining neurological integrity is well-established, with disruption correlating significantly with various forms of brain injuries, such as stroke and traumatic brain injury (Harting *et al*., 2010; Zhang, Chen and Lu, 2015; Shah and Kimberly, 2016). Among the factors affecting clinical outcomes in these conditions, the severity of edema is particularly crucial, as it can be exacerbated by alterations in BBB integrity following traumatic events (Shenaq *et al*., 2012; Wei *et al*., 2012; Shah and Kimberly, 2016). The measurement of transendothelial electrical resistance (TEER) is a key technique for assessing the integrity of BBB models *in vitro*; TEER values correlate directly with the functionality of tight junctions, which are essential for maintaining BBB properties (Kuzmanov *et al*., 2016; Henry *et al*., 2017). Beyond neuroprotection, our NRSC-EVs promoted significant recovery of BCECs barrier integrity *in vitro*, demonstrated by increased TEER values. In contrast to prior studies suggesting that NSC-derived EVs foster BCEC recovery via astrocytes (Wang et al., 2024), we observed a direct impact of NRSC-derived EVs on BCEC TEER values and viability (**Figure 5A-D**). Although not statistically significant, we also noted a trend toward decreased sodium fluorescein permeability across the endothelial monolayer upon NRSC-EV treatment (**Figure 5F**). Similar observations regarding BCEC viability have been reported previously (Deng *et al*., 2024), reinforcing the direct effect of NSC-EVs on BCECs. To our knowledge, this represents the first demonstration of a direct influence of NSC-EVs on TEER and permeability in BCECs, expanding the therapeutic potential of NRSC-EVs for enhancing BBB integrity.

The *in vivo* effects of NRSC-EVs underscore their potential for BBB recovery. In our *in vivo* model using transient middle cerebral artery occlusion (tMCAO) in rats, we observed a significantly faster resolution of brain edema in the NRSC-EV-treated group by day 7 (**Figure 6D**), with 76% of rats in this group showing complete resolution, compared to only 43% in the PBS group. Notably, while 26% of rats in the PBS group displayed persistent edema on day 7, none in the NRSC-EV group did (**Figure 6E**). This resolution of edema occurred independently of the initial size of the lesions, as the correlation between edema and lesion size observed in untreated rats was absent in animals treated with NRSC-EVs by day 7 (**Figure 6G and H**). Similarly, Webb et al. noted a reduction in edema formation in a porcine stroke model one day post-MCAO following NSC-EV treatment (Webb, Kaiser, Jurgielewicz, *et al*., 2018). In the context of brain injury, BBB impairment is a key driver of brain edema, which may manifest as cytotoxic or vasogenic edema (Harting *et al*., 2010; Shenaq *et al*., 2012; Zhang, Chen and Lu, 2015). Increased intracranial pressure due to edema correlates with BBB dysfunction, leading to significant neurological deficits and secondary neural damage (Applegate *et al*., 2016; Najjar *et al*., 2017; Vaibhav *et al*., 2020). Thus, rapid intervention to resolve edema may mitigate secondary neural damage. While therapeutic interventions like osmotherapy and decompressive craniectomy aim to alleviate these effects, they do not address the underlying molecular mechanisms promoting edema formation (Stokum *et al*., 2015; Shah and Kimberly, 2016). Our NRSC-derived EVs present an innovative approach to accelerate BBB repair and expedite edema resolution.

Despite the observed effects on edema, we noted no significant reduction in lesion size within our *in vivo* model, contrasting with earlier studies (Webb, Kaiser, Scoville, *et al*., 2018; Sun *et al*., 2019) and highlighting potential differences in experimental models or dosing regimens including timing of drug administration.

The effects attributed to EVs arise from their molecular cargo, including microRNAs and proteins that can influence recipient cell behavior, thereby facilitating paracrine signaling among cells (Yuan *et al*., 2021). Our findings indicate that cell type, rather than donor variability, is the primary determinant of EV cargo. This aligns with research by Garcia-Martin et al., which suggests that specific sequences, namely ExoMotifs and CellMotifs, dictate cell-type-specific preferences for miRNA cargo loading (Garcia-Martin *et al*., 2022). By including two hESC lines from different donors, we confirmed that donor variability minimally influences EV cargo composition, as the cargo from both hESC lines was remarkably similar (**Figure 2B and 3B**).

Among NRSC-EV miRNAs, several candidates have previously been linked to neuroprotective roles, pro-angiogenesis, and edema resolution, including *miR-21-5p*, *miR-93-5p*, *miR-92b-3p*, *miR-30a-5p*, and *miR-9-5p* (Khoshnam *et al*., 2017; Xu *et al*., 2018), in addition to various members of the *miR-17∼92* cluster, which regulate neurogenesis and NSC proliferation (Xia, Wang and Zheng, 2022). Moreover, NRSC-EVs exhibited significantly upregulated levels of *miR-145-5p* and *miR-132-3p*, both found to attenuate edema in tMCAO rodent models (Wan *et al*., 2018; Zuo *et al*., 2019). While these miRNAs may contribute to the outcomes of our study, they represent only a fraction of the total transcriptome. Further investigations are needed to understand the mechanisms underlying the EV transcriptomic cargo.

The proteomic cargo in NRSC-EVs was distinctively enriched with proteins involved in complement activation, such as C2, CFI, and CFB (**Figure 3D**). Although complement proteins are typically synthesized in the liver, the brain can produce all components of the complement system due to BBB restrictions (Brennan *et al*., 2012), which may explain their detection in EVs secreted by NRSCs. The effects of these proteins in the brain are multifaceted. Complement system activation can be neuroprotective by enhancing cytoskeletal metabolism and neurotrophic factor production, eliminating inappropriate synaptic connections, and inhibiting apoptotic pathways (Osaka *et al*., 1999; Stevens *et al*., 2007; Benoit and Tenner, 2011). However, over-activation of the complement system in the acutely injured brain can compromise neuronal integrity and exacerbate neuropathology (Brennan *et al*., 2012).

Catalase (CAT) was exclusively detected in EVs derived from NRSCs (**Suppl. Figure 4A**). Notably, previous studies have demonstrated the neuroprotective capacity of CAT in contexts of brain ischemia and neuroinflammation, where elevated levels of reactive oxygen species (ROS) can lead to neuronal cell death and dysfunction (Ali, Rakhunde and Saher, 2014). Research by Bodart-Santos et al. indicated that extracellular vesicles derived from human mesenchymal stem cells confer neuroprotective properties against oxidative stress and synaptic damage mediated by amyloid-β oligomers, a hallmark of Alzheimer’s disease. Their beneficial effects were found to depend on CAT activity, reinforcing the notion that sEVs can serve as carriers of neuroprotective components, including the CAT enzyme (Bodart-Santos *et al*., 2019). Moreover, CAT loaded into macrophage-derived sEVs has been shown to reduce oxidative stress and promote neuronal survival *in vitro* and *in vivo* in a Parkinson’s disease mouse model (Haney *et al*., 2015), and sEVs derived from rat mesenchymal stem cells were reported to attenuate oxidative stress, with the antioxidant action attributed to CAT presence in the EVs (De Godoy *et al*., 2018). These findings underscore the protective role of CAT in ischemic brain injury, where oxidative stress exacerbates cellular damage and neuronal death. The absence of CAT in sEVs derived from CMPCs and hESCs may partially elucidate the superior neuroprotective effect observed with NRSC-derived EVs in our *in vitro* models. Given the prominent role of oxidative stress in neurodegenerative processes, enhancing the activity or availability of CAT may offer a promising strategy for future neuroprotective therapies.

While this study underscores the significant therapeutic potential of NRSC-EVs, further research is essential to fully elucidate their mechanisms of action. Additionally, further investigations are needed to identify specific cargo elements driving these effects and across diverse types of brain injuries.

In summary, this data highlights the significant potential of NRSC-EVs in brain injury associated with edema formation. NRSC-EVs improved neuronal viability in an *in vitro* ischemia model, enhanced BCEC barrier integrity through increased TEER and reduced permeability, and accelerated edema resolution in the tMCAO rat stroke model. These effects can be attributed to the specific cargo of NRSC-EVs, which proved to be cell-type-specific and independent of donor variability, underscoring their unique therapeutic potential. These findings advocate for further exploration of NRSC-EVs as a promising tool for managing neurological injuries associated with edema formation, paving the way for future studies aimed at optimizing their clinical application across various brain injury contexts.

## Acknowledgements

The authors would like to express their gratitude to Maria Thaysen, Alberte Bay Villekjær Pedersen, Line Toft Ørum, Snæfríður Gunnarsdóttir, and Ann-Sophie Christiansen for their various contributions to the experiments related to the BCECs. We extend our sincere thanks to Maria Ramshøj Jensen, Lin Chen, and Merlin Till Witte for their invaluable assistance with cell culture and the collection of supernatant media. Additionally, we would like to acknowledge Charlotte Berthelsen and Sara Vitoria Brinas for their technical help.

We are also grateful to TAmiRNA for their help in conducting transcriptomic sample analysis. Biogenity has contributed to performing the proteomic data analysis, including the related figures. Technical University of Denmark has contributed to performing the proteomic sample analysis. Their expertise was essential to the successful completion of this research. Lastly, we would like to thank Charles River for their expertise and support in conducting the *in vivo* animal studies, which played a crucial role in this study.

## Methods and materials

### NRSC culture

NES^+^ NRSCs (**Suppl. Figure 9A**) were cultured as described previously (Frazier *et al*., 2025), however briefly described here. NRSCs were thawed in a ThawStar (Biolife Solutions), washed in DMEM/F12 GlutaMAX supplement (Gibco: 31331-028), supplemented with 0,2% penicillin streptomycin (Gibco: 15140-122), 1% N2 supplement CTS (GibcoTM: A1370701) and Y-27632 (FujiFilm: 257-00614) and centrifuged for 3 min at 300 g. After centrifugation, the cells were resuspended in proper volumes of DMEM/F12 GlutaMAX supplement (Gibco: 31331-028), supplemented with 0,2% penicillin streptomycin (Gibco: 15140-122), 1% N2 supplement CTS (GibcoTM: A1370701), 1‰ B27 supplement (GibcoTM: A33535-01), 10 µg/L EGF (R&D systems: 236-EG) and 10 µg/L bFGF (PeproTech: 100-18B) and Y-27632 (FujiFilm: 257-00614) and seeded on laminin-coated plates (Sigma: L2020) at a density of 500 000 cells/cm^2^. The next day, Y-27632 was removed from the media composition, in a full media change. The cells were cultured for 3-4 days until neural rosettes were visible and abundant, where media for EV isolation was collected on day 2-4 when Y-27632 was not present in the medium. The cells were continuously passaged every 3-4 days by the following procedure. The cells were dissociated for 3 min using TrypLE^TM^ select (Gibco^TM^: 12563011) until they loosened from the bottom of the well. Defined trypsin inhibitor (Gibco^TM^: R007100) was added and the cells were pipetted a few times to break them into single cell suspension. They were washed, centrifuged, resuspended and seeded at a density of 500 000 cells/cm^2^ as described above for thawing.

### CMPC culture

Clusters of CMPCs were generated as described in a previously published protocol (Kobayashi *et al*., 2024) with slight modifications for small bioreactors. HPSCs were differentiated in a 1L stirred tank bioreactor in chemically defined StemFit media (Ajinomoto Co., Inc.). They were furthermore purified using the animal-free chemically defined as described previously (Kobayashi *et al*., 2024). On day 15, the medium for EV isolation was harvested.

### hESC culture

The studies outlined in this manuscript were conducted in accordance with ethical guidelines and are covered by the following Danish ethical permits: H-20056277, H-23039171 and H-18016784 (De Videnskabsetiske Komiteer for Region Hovedstaden) for forebrain neural differentiation. hESC-A (**suppl. Figure 9B**) (NOVOe001-A) was thawed in a ThawStar (Biolife Solutions) and transferred to a wash of Nutristem® (Sartorius: 05-100-1A) supplemented with 0,2% penicillin streptomycin (Gibco: 15140-122) and Y-27632 (FujiFilm: 257-00614). They were centrifuged for 3 min at 350 g, resuspended in proper volumes of the same media and seeded in laminin-coated (Biolamina: LN521-05) T-flasks at densities of 150 000-350 000 cells/cm^2^. The next day the media was changed completely to Nutristem® (Sartorius: 05-100-1A) supplemented with 0,2% penicillin streptomycin (Gibco: 15140-122). The hESCs were cultured for 3-4 days, where media for EV isolation was collected and then passaged before reaching 70-90% confluency. To passage the cells, we would first wash them in PBS-/- (Gibco^TM^: 14190-094) and add TrypLE (Gibco^TM^: 12563011) for 5-7 min to dissociate them. We removed the TrypLE and immediately resuspended them in proper volumes of Nutristem® (Sartorius: 05-100-1A) supplemented with 0,2% penicillin streptomycin (Gibco: 15140-122) and Y-27632 (FujiFilm: 257-00614). The cells were seeded as described above and media was collected for EV isolation at day 2-4 when Y-27632 was not in the media. hESC-B (E1C3 NN GMP0050E1C3) were thawed and cultured in a similar manner.

### Immunocytochemistry

To fix cells for ICC, we firstly washed cells in PBS+/+ (Gibco: 14040-091) and then fixed them for 20 min at 4°C in 4% PFA (Thermo Scientific: J61899). After fixation, the cells were washed twice in PBS+/+, permeabilized with 0,5% Tween-20 (VWR chemicals: 663684B) and finally blocked PBS+/+ supplemented with 0,3% Triton X-100 (J.T. Baker: 9036-19-5) and 5% Donkey serum (Jackson Labs: 017-000-121). Primary antibodies were added to cells overnight at 4°C – find details in table below. The primary antibodies were subsequently removed, the cells were washed twice with PBS+/+ and secondary antibodies were added for 45 min at room temperature – find details in table below. Images were captured with a BioTek Cytation C10 Confocal imaging reader imager software V3.15.

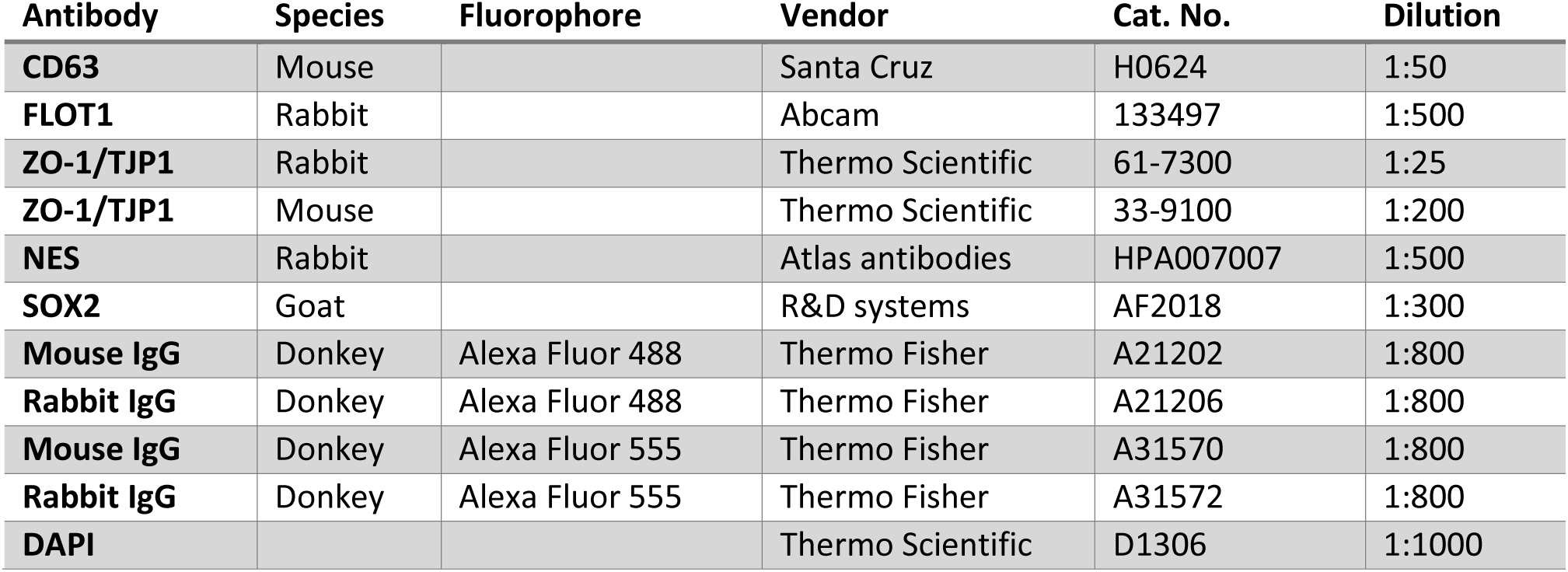

### scRNA-sequencing

scRNA-seq were performed on NRSCs as described previously (Frazier *et al*., 2025), however briefly: The 10x Genomics Chromium Single Cell 3’ Reagent kit Version 3.1 workflow was utilized, with libraries prepared with unique molecular barcoding, amplified and sequenced by Illumina NovaSeq6000. Cell Ranger pipeline was used for processing (V7.1, 10x genomics) and reads were aligned to reference genome GRCh38 (Ensembl release 107). Cells were filtered and subjected to quality control measures before visualization through UMAP was achieved in R.

### sEV isolation

sEVs were isolated from NRSCs, CMs, hESC-A and hESC-B using SEC column (IZON:IC10-35). Cell culture medium harvested after exposure to cells for 24 +/−4 h and immediately centrifuged for 10 min at 1500 g to remove cell debris. The supernatant was collected and frozen for further processing. When thawed, we centrifuged the supernatant for 10 min at 10 000 g and concentrated it with Amicon® Ultra-15 centrifugal filters 100K cutoff (Merck Millipore: UFC9100). We passed the concentrate through at a qEV10 −35 SEC column (IZON: IC10-35) with filtered (0,2 µm) PBS (Gibco: 14040-091) and thereby separated the particles depending on their size. Fractions were collected and stored at −80°C.

NRSC-EVs were also isolated via ultracentrifugation for *in vivo* experiments and *in vitro* experiments using alternative neurons. For this purpose, the collected cell culture supernatant was centrifuged for 20 min at 3000 g to remove debris and larger vesicles. The supernatant was collected and centrifuged again for 2 h at 100 000 g. The supernatant was removed hereafter and the pelleted sEVs were resuspended in PBS.

### Nanoparticle tracking analysis

The isolated EVs were characterized by concentration, size distribution and protein content. Concentration and size distribution was measured by NTA, where single particle analysis was performed using a ZetaView NTA machine (Particle Metrix) in scatter mode. The instrument was set up in accordance with the manufacturer’s recommendations and calibrated with alignment suspension containing 100 nm beads. Sensitivity was set to 80 and shutter to 100. The temperature kept constant at 23 °C. Each sample was measured in triplicates. Particle concentration and average size were calculated from the 11-position table.

### Protein quantification

Pierce^TM^ BCA Protein Assay Kit was used to determine total protein concentration. The sEV samples were diluted 1:4 in Pierce^TM^ RIPA buffer (Thermo scientific: 89900) to dissolve the lipid-bilayers and expose proteins. BCA working reagent was prepared according to vendors guidelines for standard Microplate Procedure protocol. 25µl sample was mixed with 200 µl working reagent and placed on a shaker for 30 sec to properly combine. The solution was incubated for 30 min at 37°C and then absorbance was measured at 550 nm on a SpectraMax i3x multi-mode microplate reader (Molecular devices) and captured with SoftMax Pro V7.2. Protein concentration could now be determined based on a standard curve.

### Western blotting

Western blots were performed to assess EV marker expression. NRSCs were firstly placed on ice and washed with cold PBS -/- (Gibco) before lysing them with RIPA buffer (Thermo scientific) containing 1% HALT Protease Inhibitor cocktail (100x). The lysate was kept on ice for 30 min and then centrifuged for 30 min at 12000 g. The pellet was cryopreserved at 80°C until further processing. Both EVs and cell lysates were thawed on ice, then further diluted and denatured using Laemmli SDS sample buffer, reducing (6X) (Thermo Scientific) by boiling for 5 min at 95°C. Next, we loaded 10 µg of protein onto a NuPAGE™ 4-12%, Bis-Tris, 1.0 mm, Mini Protein Gel, 12-well and NuPAGE™ 10%, Bis-Tris, 1.5 mm, Mini Protein Gel, 10-well dependent on the appropriate volume needed and additionally loaded our standard molecular weight ladder Precision Plus Protein™ Kaleidoscope™ Prestained Protein Standards (10-250 kDa). Electrophoresis was run at 200V, 150 mA, and 30W for 1 hour and 5 minutes, followed by protein transfer to PVDF membranes using iBlot™ 2 Transfer Stacks and the iBlot™ 2 Gel Transfer Device (Thermo Fisher Scientific). Successful transfer was confirmed by Ponceau S solution PierceTM at room temperature for 5 minutes and imaged using AmershamTM Imager 600. Membranes were blocked with tris-buffered saline with 0,1% Tween^TM^ (Thermo Fisher Scientific) and 5% bovine serum albumin (BSA, Sigma-Aldrich) for 1 hour at room temperature and incubated overnight with primary antibodies in blocking buffer at 4°C. Hereafter, the membranes were washed thrice for 10 min with shaking and incubated with HRP-conjugated secondary antibodies in blocking buffer for 1 h. The membranes were then washed 6 times for 5 min and incubated with Pierce™ ECL Western Blotting Substrate (Thermo Fisher Scientific) at room temperature for 1 min. Proteins were imaged with Amersham^TM^ Imager 600. A detailed list of antibodies used can be found below:

| **Antibody** | **Species** | **Vendor** | **Cat. No.:** | **Dilution** |
| --- | --- | --- | --- | --- |
| **SYNTENIN-1** | Rabbit | Thermo Scientific | PA592419 | 1:1200 |
| **CD63** | Mouse | Abcam | Ab59479 | 1:1000 |
| **CD9** | Rabbit | Abcam | Ab263019 | 1:800 |
| **CD81** | Rabbit | Abcam | Ab219209 | 1:1000 |
| **Mouse IgG HRP-conjugated** | Goat anti-mouse | R&D Systems | HAF007 | 1:1500 |
| **Rabbit IgG HRP-conjugated** | Goat anti-rabbit | Sigma-Aldrich | 12-348 | 1:3000 |

### Transmission electron microscopy

TEM was used to characterize the morphology of exosomes, conducted by Shuimu Biosciences Ltd (Beijing, China). In brief, exosome suspensions (10 µL) were applied to the copper grids coated with formvar film and incubated for 2 minutes at room temperature. The grids were then stained with a 2% Uranyl Acetate solution. After removing excess staining solution, the grids were air-dried for 10 minutes at room temperature. Dried grids were loaded into a sample holder for subsequent observation under a TEM.

### miRNA analysis

The miRNeasy Mini Kit (Qiagen) was utilized to extract total RNA from 200 µL EV suspension. EVs were homogenized with 1,000 µL Qiazol. After 10 min incubation at room temperature, 200 µL of chloroform was added and samples were incubated additionally 3 min at room temperature. Hereafter, samples were centrifuged at 12,000x g for 15 min at 4 °C. 650 µL upper phase were transferred to fresh tubes and mixed with 7 µL glycogen (5 mg/mL) to enhance precipitation. A Qiacube liquid handling robot performed precipitation, binding to miRNeasy mini columns, and washing steps. Total RNA was eluted in 30 µL nuclease-free water and stored at −80 °C until further processing. When thawed, small RNA sequencing libraries were created using 8.5 µL total RNA as input. The RealSeq Biofluids kit (RealSeq Biosciences) were used according to manufacturers instructions to generate libraries. 1 µL miND® spike-in standard (TAmiRNA GmbH) was added to each sample during the first step. Adapter-ligated libraries were amplified with 20 PCR cycles. Quality control was performed using a Bioanalyzer instrument (Agilent) with a DNA 1000 chip (Agilent). An equimolar pool of all libraries was prepared and purified on a BluePippin system (Sage Science) using 3% agarose cassettes (size range: 130-160 bp). Sequencing was performed on a NextSeq2000 instrument (Illumina) with 100 bp single-end reads.

Next-generation sequencing (NGS) analysis was performed utilizing the miND® analysis pipeline(Diendorfer *et al*., 2022). Quality assessment of the data was conducted automatically and manually by fastQC v0.12 (Andrews 2010) and multiQC v1.14 (Ewels *et al*., 2016). The samples that passed quality control were adapter-trimmed and filtered for quality by cutadapt v3.3 (Martin, 2011) and filtered for a minimum length of 17 nucleotides. Reads were mapped in 2 steps using bowtie v1.3.0(Langmead *et al*., 2009) and miRDeep2 v2.0.1.2 (Friedländer *et al*., 2012). First to the genomic reference GRCh38.p12 provided by Ensembl (Zerbino *et al*., 2018) allowing for two mismatches and secondly to miRBase v22.1 (Griffiths-Jones, 2004), filtered for miRNAs of hsa only, allowing for one mismatch. Non-miRNA reads were mapped to RNAcentral v23.0 (Sweeney *et al*., 2019) and assigned to various RNA species, to allow for a general transcriptomic composition profile. To conduct statistical analysis of the preprocessed NGS data, R v4.0 and the packages pheatmap vNA, pcaMethods v1.82 and genefilter v1.72 was employed. Differential expression analysis with edgeR v3.32 (Robinson, McCarthy and Smyth, 2010) used the quasi-likelihood negative binomial generalized log-linear model functions provided by the package. To discard love abundant miRNAs and improve false discovery rate (FDR) correction, DESeq2 independent filtering method (Love, Huber and Anders, 2014) was adapted for use with edgeR. Additional NGS QC and absolute quantification of miRNAs was done using miND® spike-ins (Khamina *et al*., 2022) based on a linear regression model.

ORA was performed using the miEAA platform(Aparicio-Puerta *et al*., 2023) and based on miRTarBase (Gene ontology). The analysis was done on the RPM counts of all miRNAs>5 RPM in all 3 replicates of the NRSCs.

### Proteomic analysis

sEVs from all cell-types were dried to ∼30 μL in an Eppendorf SpeedVac concentrator and lysed in a 1:2 lysis buffer(6M GdCl, 10mM TCEP, 40mM CAA, 50mM HEPES pH8.5 Afterwards, samples were placed in a bioruptor pico sonication water bath (Diagenode) and boiled at 95°C and sonicated for on high for 5×60 sec on/30sec off. The protein concentrations were detected with the BCA Rapid Gold assay (Thermo) and 20 μg of protein was digested per sample. The samples were then diluted 1:3 in digestion buffer (10% acetonitrile in 50 mM HEPES) and incubated for 4 h at 37°C with 1:00 enzyme-to-protein ratio LysC enzyme (MS Grade, WAKO) while shaking at 750 rpm. Hereafter, samples were furhter diluted 1:10 in digestion buffer and trypsin was added at a 1:100 enzyme-to-protein ratio overnight at 37°C under the same conditions. Digestion was followed by acidification with trifluoroacetic acid (TFA), desalting with SOLAµ SPE plates (Thermo Fisher Scientific), and processed by centrifugation at 1500 rpm between each step. To equilibrate the filters, 200 µL of 1% TFA, 3% acetonitrile was applied 2 times, after which samples were loaded. The peptides were eluted into tubes using 40% acetonitrile, 0.1% formic acid and concentrated with Eppendorf SpeedVac. Hereafter they were reconstituted in 12 µL of Solution A* (2% acetonitrile, 1% TFA) for analysis and centrifuged for 10 min at 18.000 g. Finally, the supernatant was transferred to a Nanodrop to measure peptide concentration.

The peptides were loaded onto a 2cm C18 trap column (ThermoFisher: 164946), connected in-line to a 15cm C18 reverse-phase analytical column (Thermo EasySpray ES904) using 100% Buffer A (0.1% Formic acid in water) at 750bar, using the Thermo EasyLC 1200 HPLC system, and the column oven operating at 30°C. Peptides were eluted over a 70 minute gradient ranging from 10% to 60% of Buffer B (80% acetonitrile, 0.1% formic acid) at 250 nl/min, and the Orbitrap Exploris instrument (Thermo Fisher Scientific) was run in Data-independent analysis (DIA) mode with FAIMS ProTM Interface (ThermoFisher Scientific) with CV of −45 V. A resolution of 120,000 were used to acquire the full mass spectrometry (MS) spectra, with an AGC target of 300% or maximum injection time set to ‘auto’ and a scan range of 400–1000 m/z. The MS2 spectra were obtained in DIA mode in the orbitrap operating at a resolution of 60,000 with an AGC target 1000% or maximum injection time set to ‘auto’, a normalised HCD collision energy of 32%. The isolation window was set to 6 m/z with a 1 m/z overlap and window placement on. Each DIA experiment covered a range of 200 m/z resulting in three DIA experiments (400-600 m/z, 600-800 m/z and 800-1000 m/z). Between the DIA experiments a full MS scan is performed. MS performance was verified for consistency by running complex cell lysate quality control standards, and chromatography was monitored to check for reproducibility.

SpectronautTM (version 19.1) were used to analyze the raw files and were matched against the Homo sapiens database (Uniprot proteome ID 5640). Dynamic modifications were set as Oxidation (M) and Acetyl on protein N-termini. Cysteine carbamidomethyl was set as a static modification. All results were filtered to a 1% FDR, and protein quantitation done on the MS1 level. The protein groups were inferred by IDPicker.

### *In-vitro* Neuronal stroke model

Hypothalamic-like neurons (HLNs) derived from NSCs were generated as described previously (Frazier *et al*., 2025). The cells were thawed and plated onto 96-well plates coated with 0,01 mg/ml PDL (Sigma-Aldrich: A-003-E) and 1-2 mg/ml mouse Laminin (Merck: L2020) at a density of 150.000 cells/cm^2^ and matured for 3 days in DMEM/F12 GlutaMAX supplement (Gibco: 10565018), supplemented with 1% penicillin streptomycin (Gibco: 15140122), 1% N2 supplement (Gibco: A1370701), 1% B27 Supplement (Gibco: A33535-01), 20 ng/ml GDNF (R&D systems: 212-GD) and 20 ng/mL BDNF (Sigma-Aldrich: SRP3014). After 3 days of maturation the neurons were subjected to oxygen deprivation. We utilized a two-enzyme system to deprive neurons of oxygen and by such mimic the ischemic conditions of a stroke. Glucose oxidase (Sigma: G7141) was prepared in sodium acetate buffer, while Catalase (Sigma: C1345) was prepared in potassium phosphate buffer. Various concentrations of the enzymes were tested for their capacity to deplete the medium of oxygen, and finally, a concentration of 3 U/ml glucose oxidase and 180 U/ml Catalase was selected for further experimentation. The enzymes were spiked into culture medium of DMEM/F12 GlutaMAX^TM^ supplement (Gibco: 10565018), supplemented with 1% penicillin streptomycin (Gibco: 15140122), 1% N2 supplement CTS^TM^ (Gibco: A1370701), 1% B27 Supplement (Gibco: A33535-01). 10 mmol/L Glucose were spiked into the culture at the beginning of an experiment and every 2 h to counteract the glucose consumption of glucose oxidase. Glucose- and oxygen levels were assessed using a RAPIDPoint® 500 (Siemens Healthineers), with detection ranges of mmHg pO2: 10-700 and mmol/L glucose: 1,1-41,6. At the end of the oxygen deprivation we washed the cells 3 times with PBS+/+ (Gibco: 14040-091) before fresh culture medium containing either 100 µg/ml EVs or PBS and 40 U/ml catalase were added to the neurons. The cells were then allowed to recover for 48h, before viability assays were conducted.

### *In-vitro* BBB stroke model

BCEC isolations were performed according to the Danish National Council for Animal Welfare under the license number: 2021-15-0201-01030. BCECs were isolated from male C57BL/6JRccHsd mice at 4 weeks of age. Cortices were harvested from the sacrificed animals by macroscopic dissection to remove cerebellum, white matter and meninges. The cortices were homogenized by 10 strokes with a Dounce tissue grinder (Sigma: D9188), mixed with 16% dextran (Sigma: D8821) and centrifuged for 20 min at 2500 g to separate the components of the mixture and allow capillaries to collect at the bottom. Hereafter, the capillaries were washed and resuspended them in DMEM-high glucose (Sigma D-5796) supplemented with 10% FBS (HyClone – SH30088.03), 1% NEAA (ThermoFischer – 11140050) and 1% PenStrep (Sigma – P0781). The capillary solution was subsequently filtered through a 40 µm nylon filter (Millipore: NY4104700) to collect the capillaries on top. The isolated capillaries were washed and digested with a mixture of Trypsin (Worthington: LS003703), DNAse I (Worthington: LS002139) and Collagenase III (Worthington: LS004182) for 60 min before they were washed and seeded onto a T25 coated with collagen IV (Sigma – C5533) and fibronectin (Sigma – F1056). After allowing the capillary fragments to attach for 2-3 h, the media was changed to DMEM-high glucose (Sigma D-5796) supplemented with 10% FBS (HyClone – SH30088.03), 1% NEAA (ThermoFischer – 11140050) and 1% PenStrep (Sigma – P0781) and 4 ug/ml Puromycin (Sigma – P9620). The puromycin was added to kill contaminating pericytes and glial cells while allowing endothelial cells to survive, due to their expression of efflux transporter P-glycoprotein. Media change was performed after 2 days in culture where puromycin was removed and after 2 additional days endothelial cells could be plated onto transwell inserts. To induce stroke-like conditions of starvation and waste accumulation, the media was omitted from the basolateral side of the transwells.

### Cell viability determination

To determine the viability of BCEC CellTiter-Glo® Luminescent Cell Viability Assay (Promega: G7570) was used. The same assay was used to determine neuronal cell viability, multiplexed with RealTime-Glo^TM^ MT Cell Viability Assay (Promega: G9711).

An endpoint assay for RealTime-Glo^TM^ Cell Viability Assay was employed to assess cell viability using reducing potential as an indicator of metabolically active cells. 1:1000 of MT Cell Viability Substrate and 1:1000 NanoLuc® Enzyme was added were added to culture medium consisting of DMEM/F12 GlutaMAX^TM^ supplement (Gibco: 31331-028), supplemented with 0,2% penicillin streptomycin (Gibco: 15140-122) and 1% N2 supplement CTS (GibcoTM: A1370701). After vortexing, the mixture was equilibrated to 37°C and added to the neurons that were hereafter incubated for 30 min at 37°C. Following incubation, the luminescence was determined on a SpectraMax i3x multi-mode microplate reader (Molecular devices) and captured with SoftMax Pro V7.2 and percentage conversion was based on normalization to healthy control wells that had not been subjected to oxygen deprivation.

CellTiter-Glo® Luminescent Cell Viability Assay was utilized to assess cell viability by total ATP content as an indicator of metabolically active cells. An equal volume of CellTiter-Glo® Reagent was added to the culture medium and mixed well on an orbital shaker to induce lysis. After allowing the mixture to rest for 10 min, we determined the luminescence with a SpectraMax i3x multi-mode microplate reader (Molecular devices) and captured with SoftMax Pro V7.2 for neurons and CLARIOstar (BMG Labtech) for BCEC. Percentage conversion was also conducted by normalization to healthy control wells that had not been subjected to oxygen deprivation.

A few technical replicates (i.e. wells) were excluded from the analysis due to significant deviations from the expected value ranges, which could not be attributed to experimental conditions. A two-way ANOVA was chosen for analysis as we have >3 categorical groups and 2 factors, namely the treatment factor i.e NSC-EV, CM-EV ect. and the replicate factor i.e batch differences. Post-hoc multiple comparison analysis was subsequently performed to identify the groupwise differences, here, a Dunnetts test was used where groups NSC-EV, CM-EV, hESC-A, hESC-B all were compared to control group PBS. 0H, 0,1H and 5,0H were kept as visual controls on graph, but not included in ANOVA

### Measurement of resistance (TEER)

To measure the resistance of the BCEC, transwell inserts were transferred to a CellZScope (NanoAnalytics: CellZScope+) for continuous measurements. CellZScope measurements were conducted in a spectrum of 1Hz-100kHz and captured in CellZScope software version 4.4.6 build 210520. Wells that reached higher than 50 Ω*cm^2^ within the first hour were excluded from the experiment to assure successful injury of the BCECs. TEER was measured on different occasions to obtain biological replicates. Depending on the number of technical replicates, each recording would occur at different timepoints, which meant the different recording did not share a common x-axis. To circumvent this, a common time grid was made, and the data was interpolated, to estimate what the TEER values would be at the time points in the common time grid. The following formula was used for interpolation XLOOKUP(Common time grid cell; Run time column; TEER column; ; 1) in excel (Version 2503, build 18623.20266). Paired t-test was performed on the AUCs with 95% confidence level.

For endpoint TEER measurements, a cup electrode by EVOM^TM^ TEER meter (World Precision Instruments) was employed to measure the resistance in ohm and multiplied by the surface area of the filter to determine the TEER in Ω*cm^2^.

### Permeability measurement

To assess the barrier integrity of our BCECs, an *in vitro* transport study was performed. A sodium fluorescein (Sigma-Aldrich: F6377-100g) standard solution of 5 mg/ml was added to the apical side of a transwell system, resulting in a final concentration of 500 µg/ml. The cells were then incubated for at 37°C with agitation (90 RPM), and were collected from the receiver compartment at 60, 90, 105, 120 minutes, with the volume replenished after each collection. Furthermore, samples were taken from each donor compartment at 0 and 120 minutes. All samples were transferred to a black, clear-bottom 96-well plate, together with a blank control consisting of cell culture media. The fluorescence could now be measured using a NOVOstar (BMG Labtech) at excitation at 460 nm and emission at 515 nm. The sodium fluorescein concentrations quantified by comparison to a standard curve, that was prepared fresh for each individual experiment. Total amount transported was plotted over time, with the slope representing the steady state flux (J), which was applied to calculate the apparent permeability coefficient (P_app_) using the following equation. 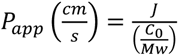, with J corresponding to the steady state flux, M_w_ representing the molecular weight of sodium fluorescein and C_0_, the donor concentration at 0 minutes.

### *In-vivo* stroke model

Animal studies were conducted in Charles River Finland and performed according to the EU Directive 2010/63/EU and were approved by the Novo Nordisk Animal Welfare Body. The *in vivo* tMCAO experiments conducted in this study were in compliance with the ethical standards as specified by the following national Finnish guidelines: 1) Act on the Protection of Animals Used for Scientific or Educational Purposes (497/2013), 2) Government Decree on the Protection of Animals Used for Scientific or Educational Purposes (564/2013), 3) License for animal experiment approved by National Animal Experiment Board [ESAVI/20200/2021]. They furthermore adhered to the following EU and international guidelines: 1) Directive 2010/63/EU, 2) Commission recommendation 2007/526/EC, 3) Guide for the Care and Use of Laboratory Animals (Guide), Eighth Edition (National Research Council 2011). Rats were inspected daily and euthanized if they met predefined humane endpoints.

96 male Wistar rats (approx. 250g) were subjected to tMCAO and of these, 10 were excluded due to insufficient lesion size. The procedure was initiated with analgesic regimen starting at least 30 min prior to start of the surgery with dosing of 0,05 mg/kg buprenorphine in 8-hour intervals for the first day and 12-hour intervals for additionally 2 days. On day 0 the rats were anesthetized (2-5% isoflurane for induction, 1.5-2.5% isoflurane for maintenance) and placed on their back on a 37°C heating pad with rectal probe monitoring. Ringer Lactate was administered (s.c.) to prevent dehydration, and lidocaine was injected in the skin before performing a neck incision under antiseptic conditions. After exposing the right common carotid artery (CCA), a filament with a silicone covered tip was inserted into the internal carotid artery 22-23 mm up to the origin of the middle cerebral artery to block cerebral blood flow. The filament was left in place for 90 min. The neck wound was carefully reopened after the 90 min and the filament was removed to achieve reperfusion. The wound was sutured with a nylon thread, and the rats were closely monitored for the next 2 h for any post-surgical complications. Animals were rehydrated at day 1, 2 and 3 with subcutaneous administration of body warm Ringer Lactate (10 ml/kg) when handled during other procedures.

The rats were dosed with 114 µg NRSC-EVs (n=) or equivalent PBS volumes (n=) at 3 h, 24 h and 48 h following the reperfusion by intravenous bolus dosing through the tail vein.

To verify the success of the tMCAO and to provide an early quantification of the lesion size, DWI-MRI was performed 60 min after the insertion of the filament and prior to reperfusion. A successful lesion was confirmed by a 20% or more drop in apparent diffusion coefficient (ADC) of the ipsilateral hemisphere, compared to the contralateral hemisphere. The ADC map was obtained in 12 min by spin-echo based diffusion weighted MRI sequence with in-plane resolution of 118 mm and 0.7 mm slice thickness. Prior determined exclusion criteria entailed that rats with lesions smaller than 50 mm^3^ and larger than 220 mm^3^ was excluded from the experiment.

Absolute T2-MRI was performed on day 3, 7 and 28 to assess the development of lesion size and edema. Multi-slice multi-echo sequence was utilized with the following parameters: TR=2.5 s, 12 different echo times (10-120 ms in 10 ms steps) and 4 averages. 18 coronal slices with 1 mm thickness were acquired using field-of-view 30×30 mm^2^ and 256×128 imaging matrix. Lesion volumetric analysis was performed blinded by contrasting differences between lesioned and healthy tissues in the ipsilateral side, considering reference values and contralateral hemisphere as internal control.

Neurological and motor deficits were evaluated on day −3, 3, 7 and 28 using 3 complimentary blinded behavioral tests, namely 7-point neuroscore, 20-point neuroscore and limb-placing test. The 7-point neuroscore focused on motor and behavioral deficits, scoring contralateral forelimb placement and sensorimotor function with scores from 0-6, where 0 reflect severe deficits and 6 reflects normal behavior. The 20-point neuroscore aimed to evaluate the global neurological function, including spontaneous movements, forelimb strength and reflexes, with scores ranging from 0-20 with 0 showing severe neurological deficits and 20 showing normal function. Finally, the limb-placing test is used for evaluating sensorimotor integration via tactile and proprioceptive stimuli applied to all 4 limbs and scoring them from 0-2 with 0 indicating no response and 2 indicating normal response.

### Statistical analysis

Data were collected from at least three independent experiments and presented as mean ± standard deviation unless otherwise specified. Student’s *t*-test were used to compare two groups of variables, while ANOVA were used to compare multiple groups, followed by appropriate post-hoc tests. Statistical tests were performed in GraphPad Prism 10.4.1 software and P<0,05 were considered statistically significant.

### Use of generative AI and AI-assisted technologies

Generative AI tools were used by the authors to improve readability and language of the original text. No generative AI tools were used to create any of the data or to create or alter images.

---

**Supplementary figure 1.**
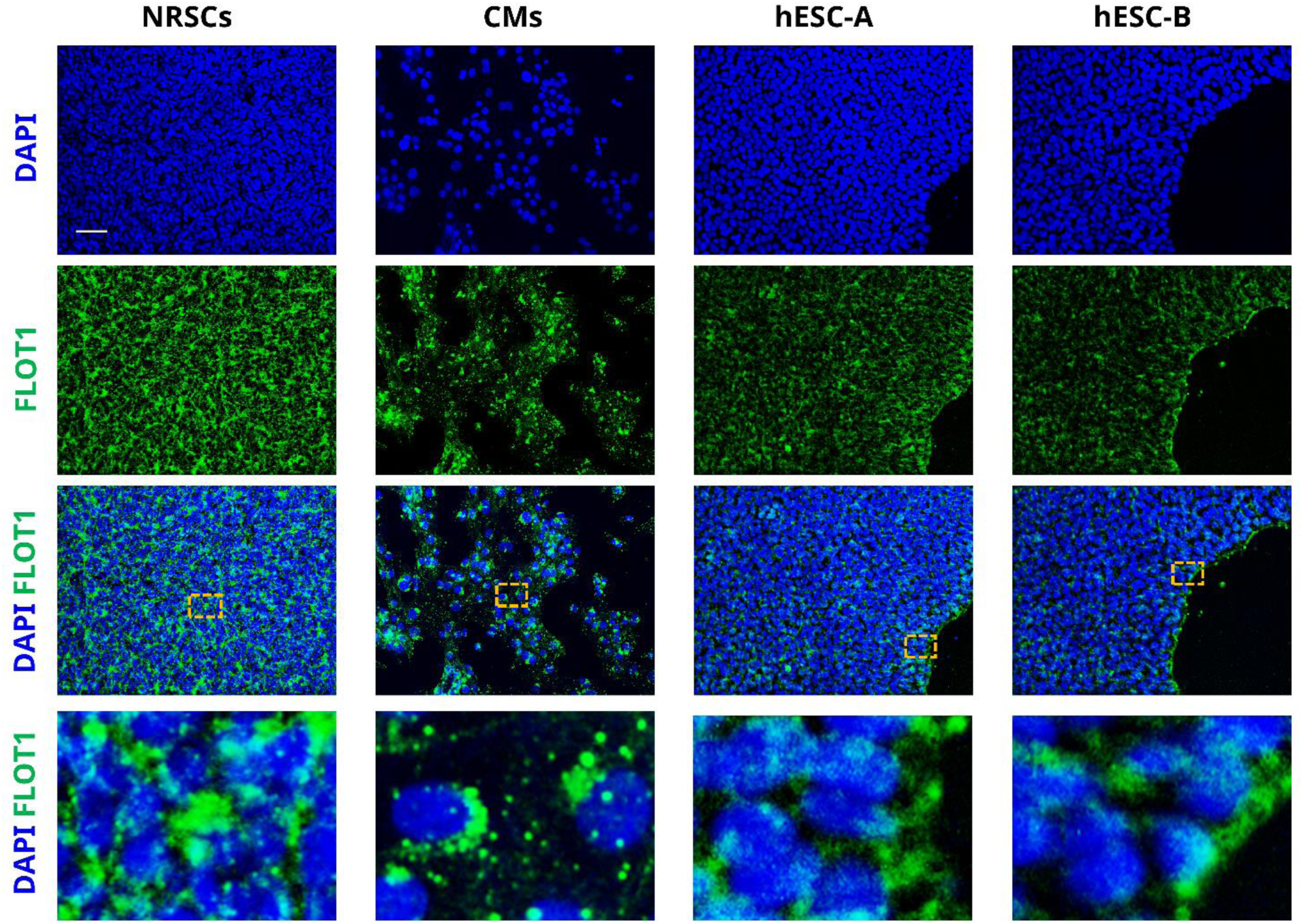
Small extracellular vesicles display cell-type specific distribution of FLOT1. Representative immunofluorescence images displaying the distribution of EV marker FLOT1 in NRSCs, CMPCs, hESC-A and hESC-B. Scalebar represents 50 µm.

**Supplementary figure 2.**
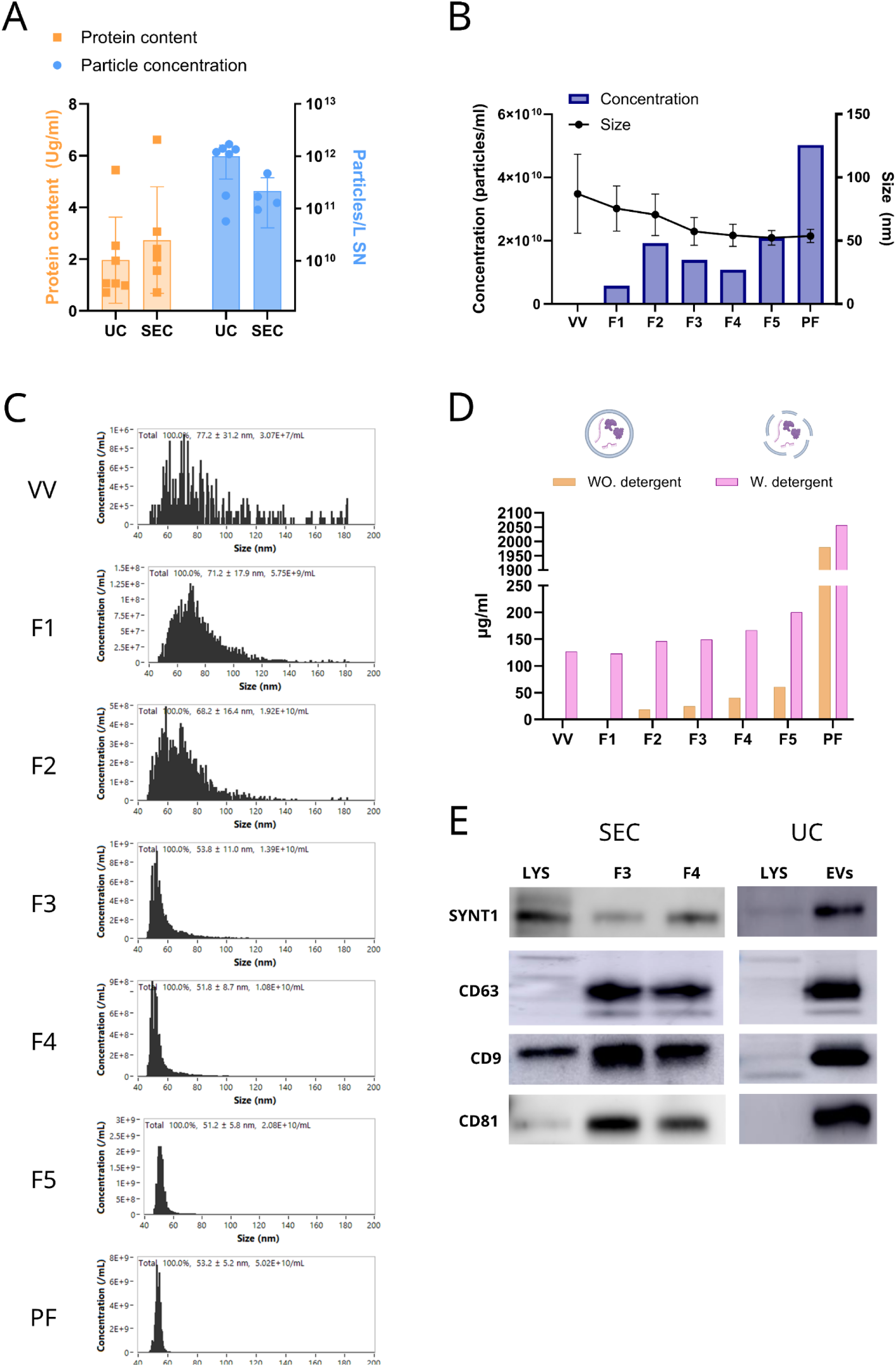
Small extracellular vesicles can be isolated from NRSC culture medium. **(A)** Collective data on sEVs isolated from NRSC culture medium by UC or SEC showing protein contents (Left y-axis) and particle concentration (right y-axis). Data shown as mean+/−SD. It should be noted that UC and SEC samples have been measured on separate instruments. N=7 for UC, N=6 for SEC protein concentration and N=4 for SEC particle concentration **(B)** Particle concentration and size distribution of different fractions isolated from SEC columns reveal that particles in the exosomal size range of 30-150 nm can be found in F1-F5. Size shown as mean+/−SD, Concentration shown as mean. N=1. **(C)** Individual size distribution profiles (raw data for B) dependent on fractions. **(D)** Proteins content of the different fractions reveals that the majority of protein in found in PF, indicating that we have low protein contamination. N=1. VV = void volume, F1=fraction 1, F2=Fraction 2, F3=Fraction 3, F4=Fraction 4, F5=Fraction 5 and PF=Protein fraction. **(E)** Western blots of NRSC-EVs isolated by SEC or UC, comparing EV marker expression in whole cell lysate (LYS) compared to EV isolates. UC=ultracentrifugation, SEC=size exclusion chromatography

**Supplementary figure 3.**
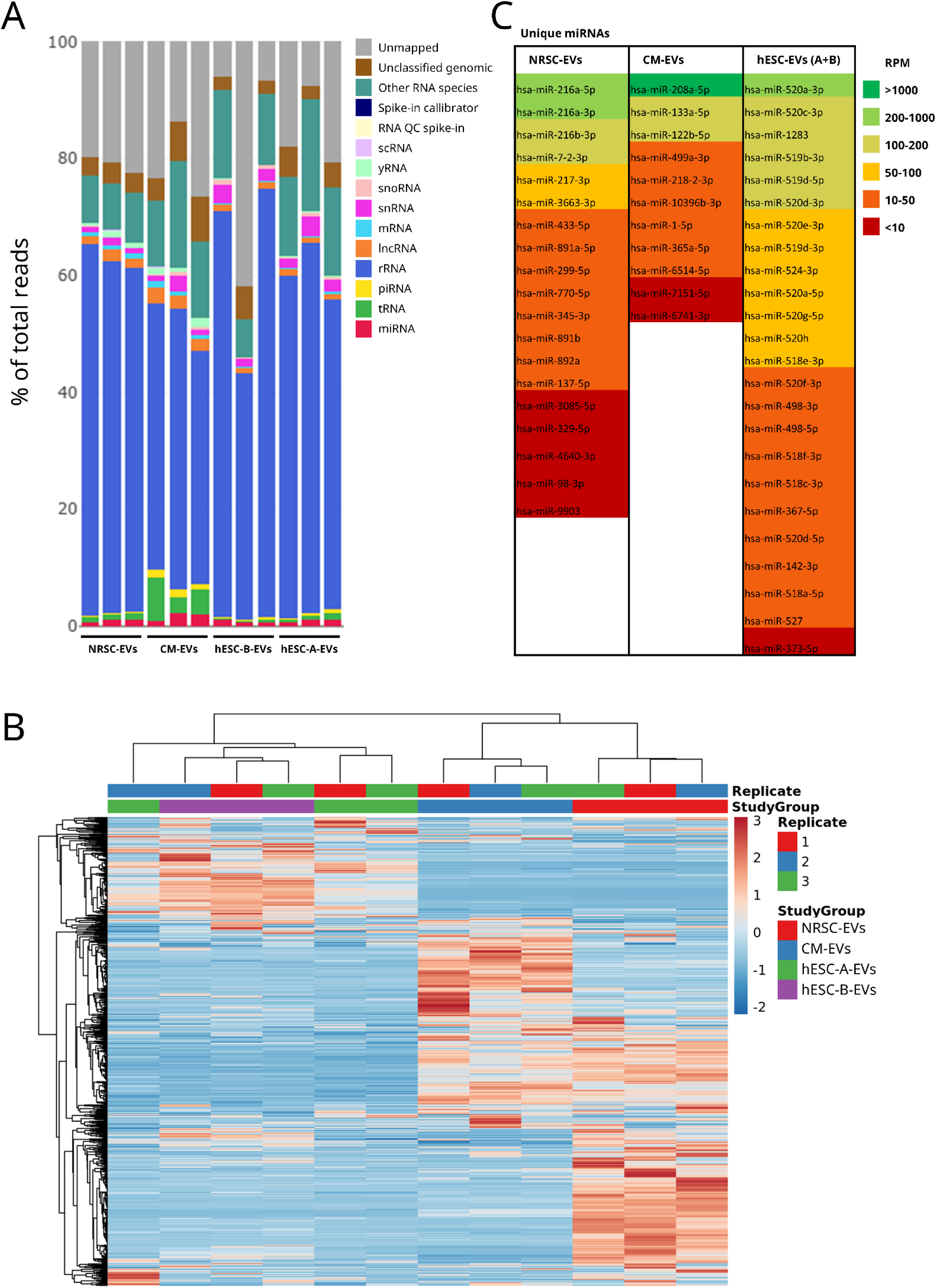
NRSC-EVs carry distinct transcriptomic profile. **(A)** Reads classification of entire EV transcriptome of the different samples shown as percentage of total reads indicating relative abundance in each sample. N=3. **(B)** Heatmap visualizing the unique patterns of the EV miRNA composition dependent on the cellular source. Heatmap color represents the RPM normalized reads in individual samples. N=3. **(C)** Unique miRNAs in each group sorted according to abundance with the most abundantly unique miRNAs at the top.

**Supplementary figure 4.**
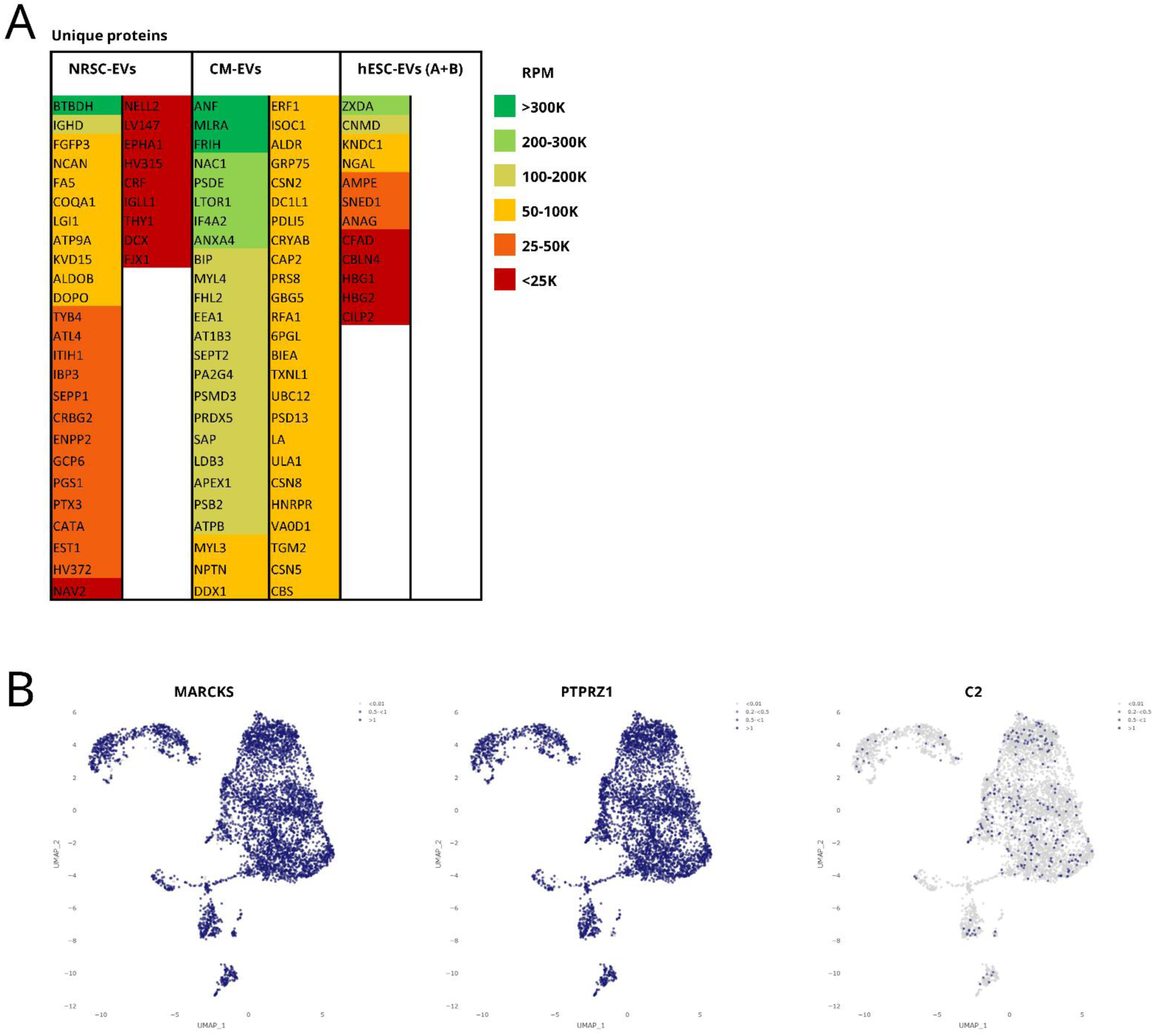
NSC-EVs carry distinct proteomic profile. **(A)** Unique proteins in each group sorted according to abundance with the most abundantly unique proteins at the top. **(B)** UMAPs based on scRNA-seq data indicating the expression levels of proteins MARCKS, PTPRZ1 and C2.

**Supplementary figure 5.**
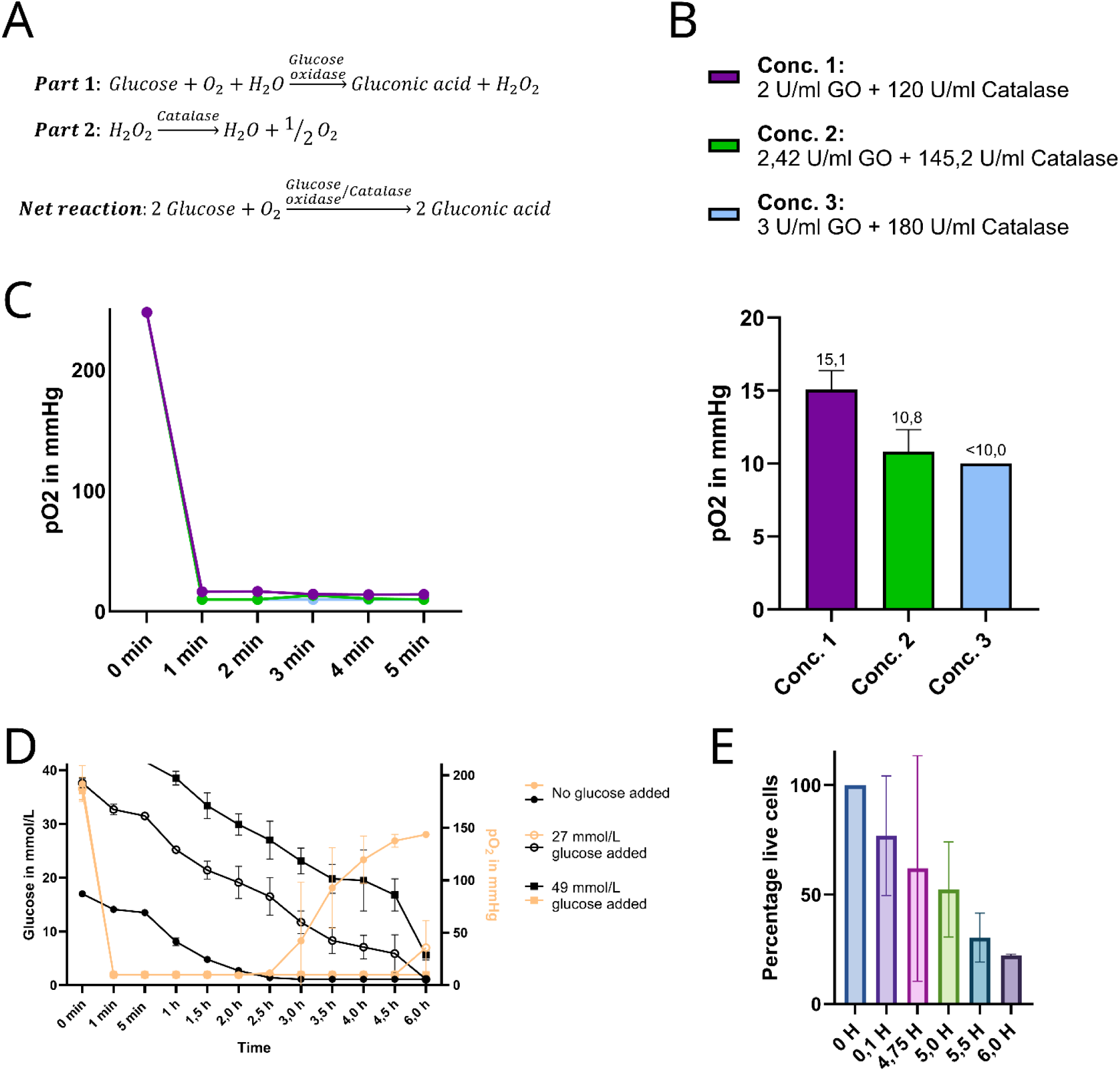
Enzymatic oxygen deprivation by glucose oxidase and catalase. **(A)** Enzymatic reaction of glucose oxidase and catalase to induce oxygen deprivation in cell culture medium. **(B)** Three concentrations of glucose oxidase and catalase were tested and the *pO*_2_ was measured to determine the optimal concentration for *O*_2_depletion. Data shown as mean +/− SD *pO*_2_within the first 5 min. n=5, N=1. **(C)** Consecutive measurements of *pO*_2_ to determine how fast the *O*_2_ was depleted from the medium. The *O*_2_ was depleted in less than 1 minute. N=1. **(D)** Different glucose concentrations were added to investigate the rate of glucose consumption in order to extend the oxygen deprivation. The glucose concentrations were tested on conc. 3 of glucose oxidase and catalase. Data is shown as mean +/− SD. N=3. **(E)** Viability measurements of neurons subjected to different durations of enzymatic oxygen deprivation. Data shown as mean +/− SD. N=2-3.

**Supplementary figure 6.**
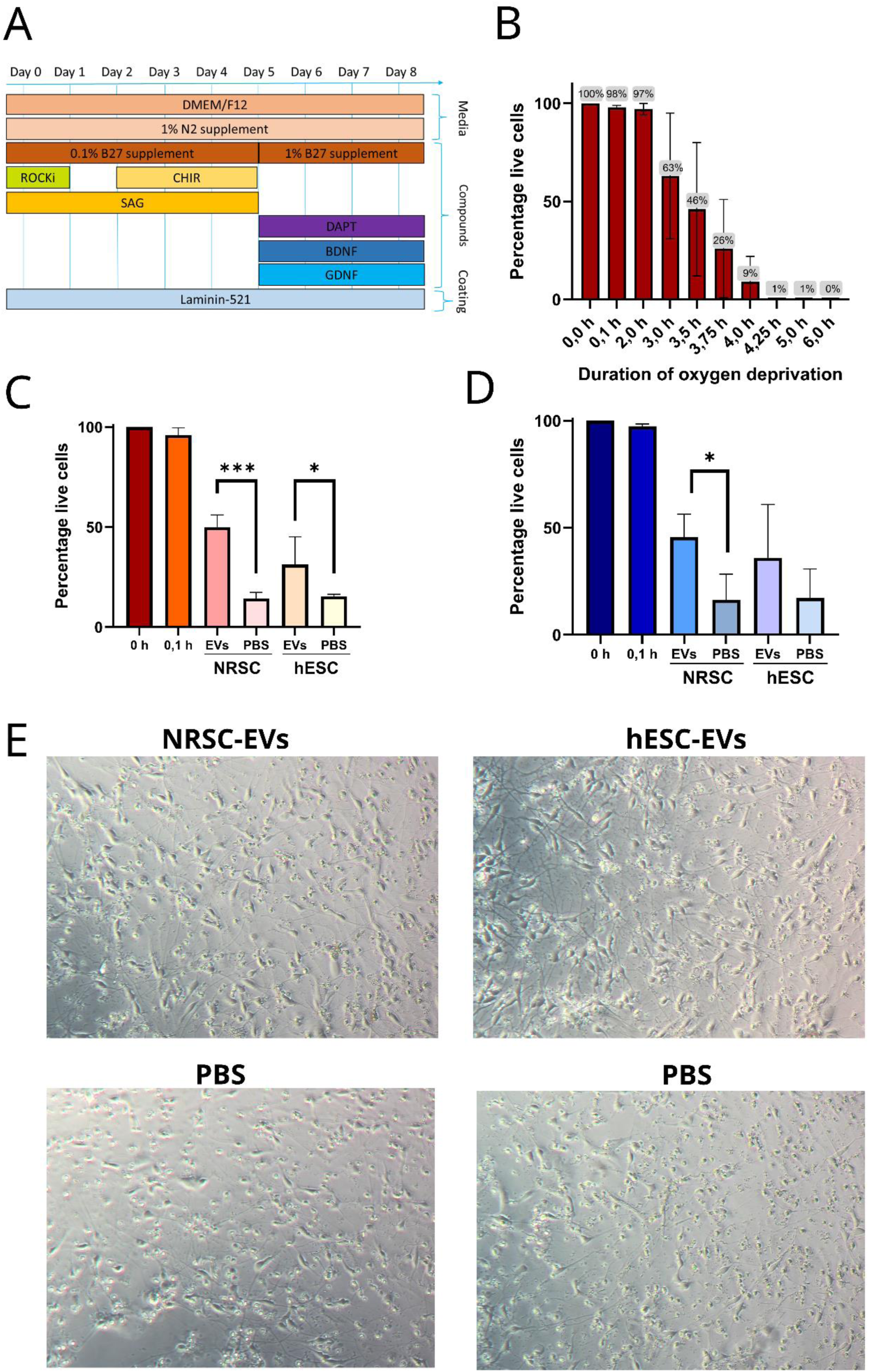
NRSC-EVs show neuroprotective effect in alternate *in-vitro* neuronal stroke model. **(A)** Schematic representation of alternative protocol for deriving neurons from NRSCs for oxygen deprivation experiments. **(B)** Viability measurements based on total ATP, for investigation of optimal duration of oxygen deprivation for a window of intervention. **(C)** Viability measurement based on reducing potential of alternate neurons after 2 days of treatment with EVs derived from NRSCs and hESC-As. **(D)** Viability measurement based on total ATP level of alternate neurons after 2 days of treatment with EVs derived from NRSCs and hESC-A. All EVs treated groups and PBS were exposed to 3,75 H of oxygen deprivation. Data is shown as mean +/− SD. Statistical tests done are unpaired t-tests, *=P<0,05 ***=P<0,001. **(E)** Brightfield images showing the morphological changes in neurons after 2 days of treatment with EVs derived from NRSCs and hESC-As. Images are taken in 20X magnification.

**Supplementary figure 7.**
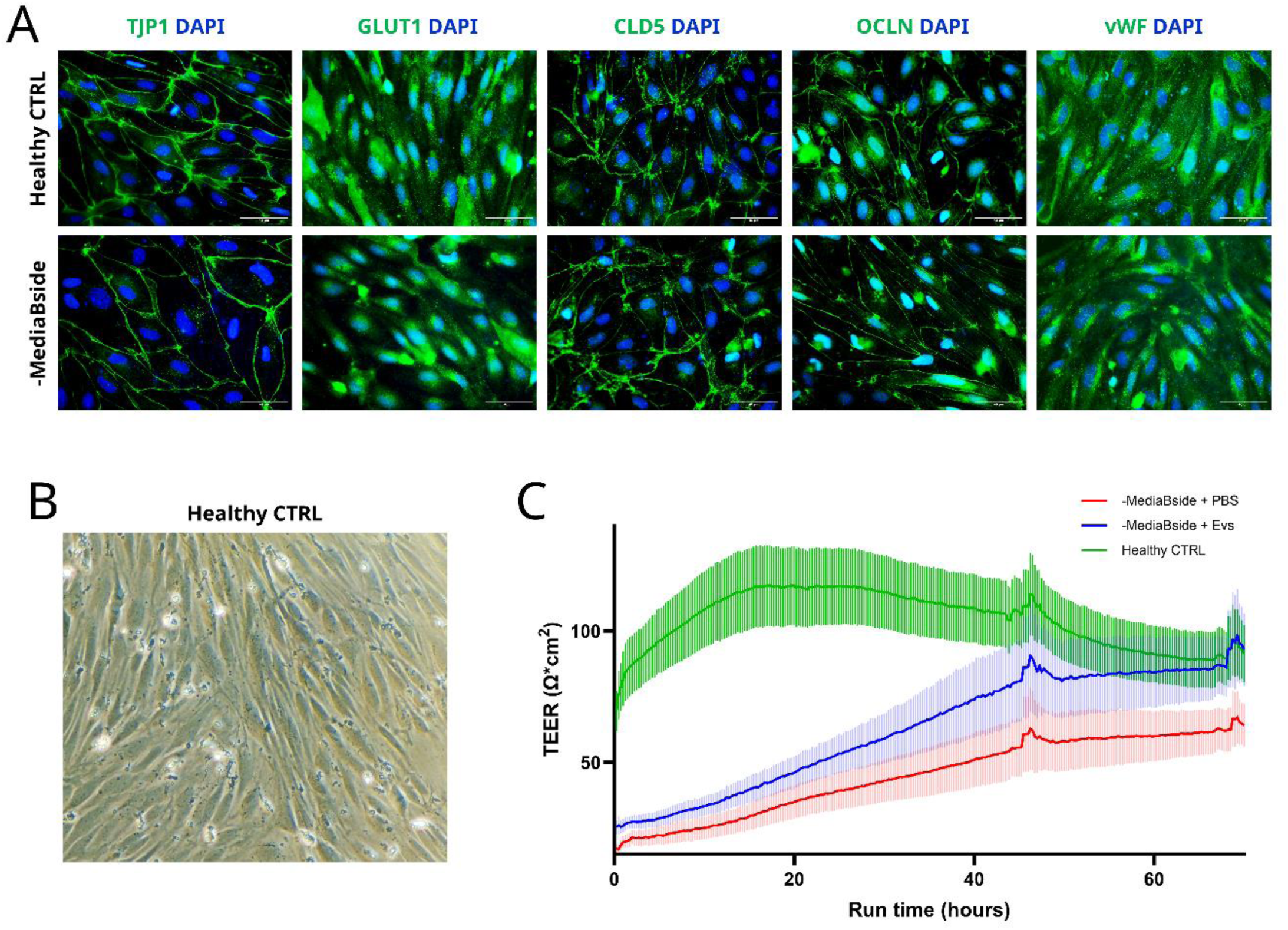
BBB model characterization and healthy controls. **(A)** Representative immunofluorescence images showing the distribution of tight junction proteins TJP1, Occludin and Claudin-5 as well as endothelial cell markers GLUT1 and vWF in BCECs on day 9 after isolation. Scalebar represents 40 µm **(B)** Brightfield image of BCEC healthy control monolayer on day 9 after isolation. Photos are taken in 20X magnification. **(C)** Raw TEER measurements before normalization of BCECs over 70 hours after addition of NRSC-EVs or PBS shown with healthy control. Data is shown as mean+/− SEM.

**Supplementary figure 8.**
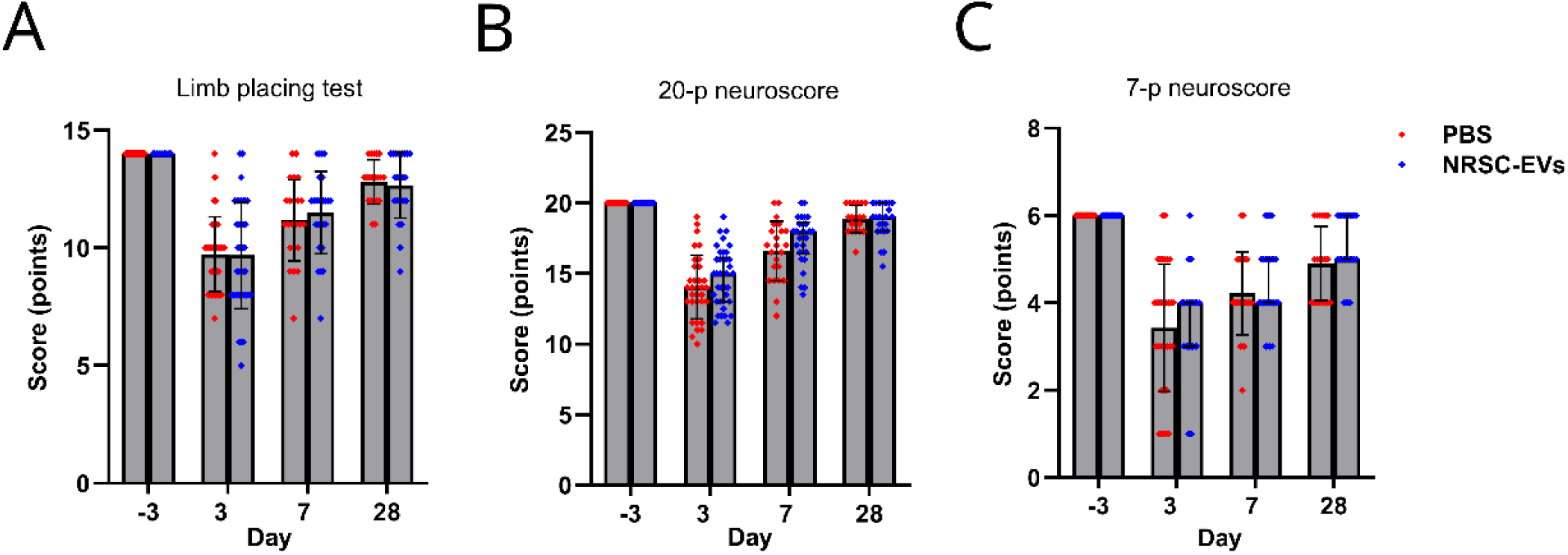
Behavioral tests of rat tMCAO model subjected to NRSC-EV treatment. **(A)** Limb placing test of rats subjected to tMCAO and treated with either NRSC-EVs or PBS. Results showed no significant differences. **(B)** 20-p neuroscore showed no significant differences, but a slight tendency toward a better performance in the NRSC-EV group. **(C)** 7-p neuroscore showed no significant differences. Data shown as mean +/− SD.

**Supplementary figure 9.**
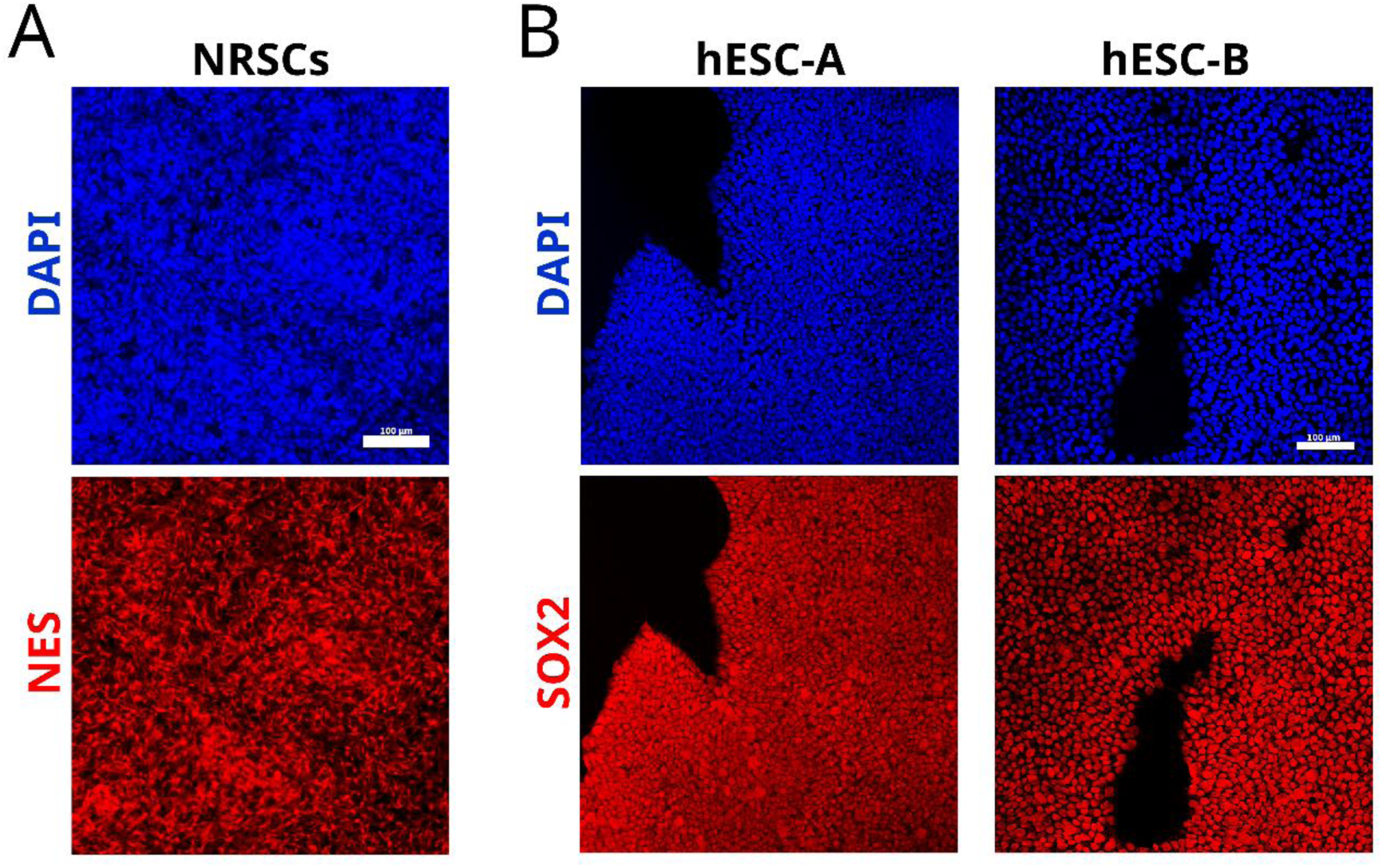
NSC and hESCs are positive for cell-type specific markers. **(A)** NRSCs showing positive for NSC-marker NES. Scalebar represents 100 µm. **(B)** Both cell lines of hESCs are positive for proliferation marker SOX2. Scalebar represents 100 µm.

